# Identification and prediction of developmental enhancers in sea urchin embryos

**DOI:** 10.1101/2021.03.26.436886

**Authors:** César Arenas-Mena, Sofija Miljovska, Edward J. Rice, Justin Gurges, Tanvi Shashikant, Sevinç Ercan, Charles G. Danko

## Abstract

**Background:** The transcription of developmental regulatory genes is often controlled by multiple cis-regulatory elements. The identification and functional characterization of distal regulatory elements remains challenging, even in tractable model organisms like sea urchins.

**Results:** We evaluate the use of chromatin accessibility, transcription and RNA Polymerase II for their ability to predict enhancer activity of genomic regions in sea urchin embryos. ATAC-seq, PRO-seq, and Pol II ChIP-seq from early and late blastula embryos are manually contrasted with experimental *cis-*regulatory analyses available in sea urchin embryos, with particular attention to common developmental regulatory elements known to have enhancer and silencer functions differentially deployed among embryonic territories. Using the three functional genomic data types, machine learning models are trained and tested to classify and quantitatively predict the enhancer activity of several hundred genomic regions previously validated with reporter constructs in *vivo*.

**Conclusions:** Overall, chromatin accessibility and transcription have substantial power for predicting enhancer activity. For promoter-overlapping cis-regulatory elements in particular, the distribution of Pol II is the best predictor of enhancer activity in blastula embryos. Furthermore, ATAC- and PRO-seq predictive value is stage dependent for the promoter-overlapping subset. This suggests that the sequence of regulatory mechanisms leading to transcriptional activation have distinct relevance at different levels of the developmental gene regulatory hierarchy deployed during embryogenesis.

## Background

Transcriptional regulatory elements (TREs) (1) are the primary drivers of differential gene expression during metazoan development (2–4). Whereas promoters are TREs easily found by association with the transcription start sites (TSSs) of genes, the identification and functional characterization of TREs distal to TSSs (enhancers and silencers) remains challenging. The oftentimes complex expression of developmental regulatory genes (that is, transcription and signaling factors) is primarily controlled by distal regulatory elements (2,5), but only a few *cis-*regulatory modules (CRMs) that constitute the essential transcriptional nodes of developmental gene regulatory networks are functionally understood (6). Histone marks such as H3K27ac (7), chromatin accessibility (8,9) or transcription initiation (10) facilitate the identification of active enhancers. In addition, high-throughput reporter assays allow genome-wide testing of enhancer activity (11). Each of these approaches has particular advantages and limitations (12), and, despite recent progress, most TREs remain largely uncharted (13,14).

Several experimental advantages have facilitated the exhaustive reconstruction of developmental gene regulatory networks (GRNs) in sea urchin embryos (15–17). The analysis of topological GRN models reveals an uneven distribution of regulatory sub-circuit motifs along the GRN hierarchy sequentially deployed during sea urchin embryogenesis (4). Accordingly, the structure and Boolean logic of the TREs serving the nodes of these sub-circuits changes during development too (16). In sea urchins, enhancer and silencer activities of TREs can be tested by lack of function in bacterial artificial chromosome (BAC) reporter constructs microinjected into zygotes (18,19), or by gain of function in much smaller plasmid reporters, which oftentimes use heterologous promoters (20). These exogenous reporters replicate along with the genome (21,22), with the much larger than plasmids BACs providing a closer approximation to the natural genomic context and more faithfully reproducing the endogenous expression. In addition, BACs maintain endogenous promoters in the context of gene-reporter translational fusions. Despite these advantages, the identification and testing of developmental TREs remains challenging due to the low throughput of existing experimental approaches.

Evolutionary sequence conservation has been routinely used for the identification of potential TREs in sea urchins (4), although conservation is not informative about the stage of which regulatory activity, and the obscure cause of diverse evolutionary rates for regulatory sequences (23,24) raises great uncertainty regarding false positive and negative rates. As in other model systems, chromatin accessibility also facilitates the identification of candidate enhancers in sea urchin embryos (19,25). In addition, a parallel reporter method enabled the enhancer activity test of several hundred CRMs associated with 37 developmental regulatory genes during sea urchin development (26,27). We used this functional quantification to train and test machine learning model predictors of developmental enhancer activity from various genomic profiles: chromatin accessibility estimated by ATAC-seq (28), RNA Polymerase II (Pol II) distribution detected by ChIP-seq, and transcription initiation estimated by the analysis of the transcription run-on assay PRO-seq (29), which detects the location of paused and elongating RNA Pol II at base pair resolution (**Fig. 1 A**). Our analysis reveals that chromatin accessibility and transcription both enable enhancer activity prediction, and that the predictive power of these genomic profiles declines during development for the subset of promoter proximal TREs, suggesting a sequence of regulatory shifts at different levels of the gene regulatory hierarchy that is deployed during development.

**Figure 1.**
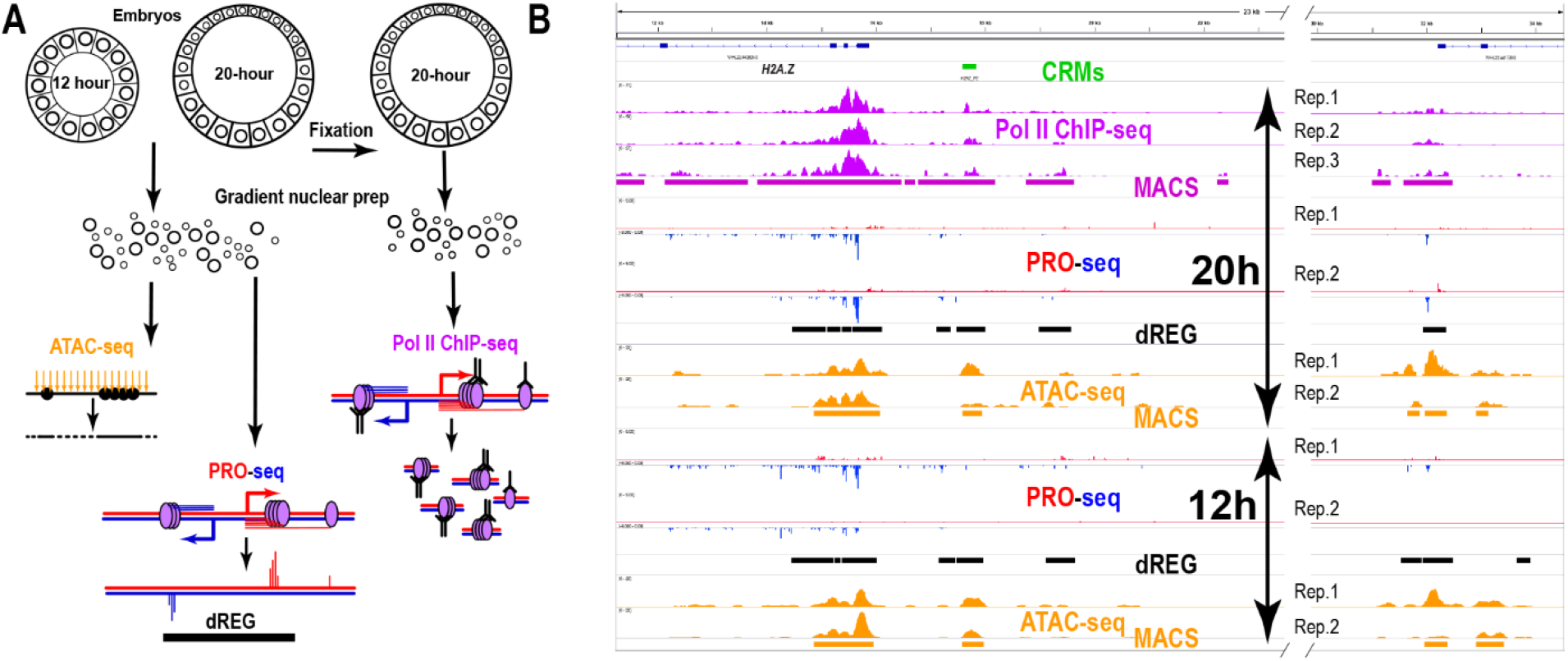
ATAC-seq, PRO-seq and Pol II ChIP-seq are used for the identification of TREs. - A, experimental outlines of the 3 genomic profiles used.
- B, IGV browser snapshot of replicate genomic profiles at the *H2A.Z* locus, a highly expressed gene (33), left, which also includes a gene expressed at lower levels, right side. Number of 3’ end reads per million of PRO-seq run-on transcripts are shown for the plus and minus strands. PRO-seq peaks mark transcriptional pause sites. MACS peak and dREG TRE predictions for the combined data sets are shown underscoring each genomic profile. The CRM panel underscores a genomic region with enhancer activity tested by deletion in large reporter constructs (Hajdu et al., 2016). PRO-seq and ATAC-seq profiles are set to the same scale between 12 and 20 hour stages, with the range indicated between brackets at the beginning of each track.

## Results

### Genomic distribution of chromatin accessibility, Pol II and transcription

Chromatin accessibility and transcription were used for the identification of candidate developmental enhancers in 12 and 20 hour sea urchin embryos. The genomic profile of 3’ end transcripts identified by PRO-seq was analyzed with dREG (**Fig. 1 A**), a support vector regression tool trained to identify TREs associated with active chromatin marks using the shape of transcription (10,30). In addition, we mapped chromatin accessibility with ATAC-seq in both stages, and in 20 hour embryos the distribution of RNA Pol II using ChIP-seq (**Fig. 1 A**), which has been also associated with active enhancers (31). In 12 and 20 hour embryos, dREG identified 43,912 and 56,753 TRE predictions or “peaks”, respectively, while a total of 238,838 and 258,515 ATAC-seq peaks, respectively, were called by MACS (**QC reports in supplementary information**). In 20 hour embryos, 554,846 Pol II ChIP-seq peaks were called.

The dynamic range of the read distribution at peak calls expands several orders of magnitude for the three functional genomic data types (**Fig. 2 B and D**). The Pearson correlation of total PRO-seq reads at dREG peak calls of biological replicates is higher for 12 hour embryos (R = 0.88, p-value < 2.2 ×10^−16^, Spearman = 0.71, **Fig. 2 A**) than for 20 hour embryos (R = 0.35, p-value < 2.2 ×10^−16^, Spearman= 0.75, **Fig. 2 C**). Similarly, higher correlation for ATAC-seq profiles at MACS peak calls is found for 12h embryos (R = 0.91, p-value < 2.2 ×10^−16^, Spearman= 0.79, **Fig. 2 A**) relative to 20 hour embryos (R = 0.62, p-value < 2.2 ×10^−16^, Spearman= 0.46, **Fig. 2 C**). The low correlation of PRO-seq biological replicates may be due in part to inherent batch heterogeneity associated with seasonal and genetic variability in the wild populations from which the embryos where obtained. This natural variation may shift the relative timing of major transcriptional regulatory changes during and prior to the 20 hour embryo stage, as previously reported (32,33), along with the associated histone modification and chromatin accessibility signals (**Fig. 2**).

**Figure 2.**
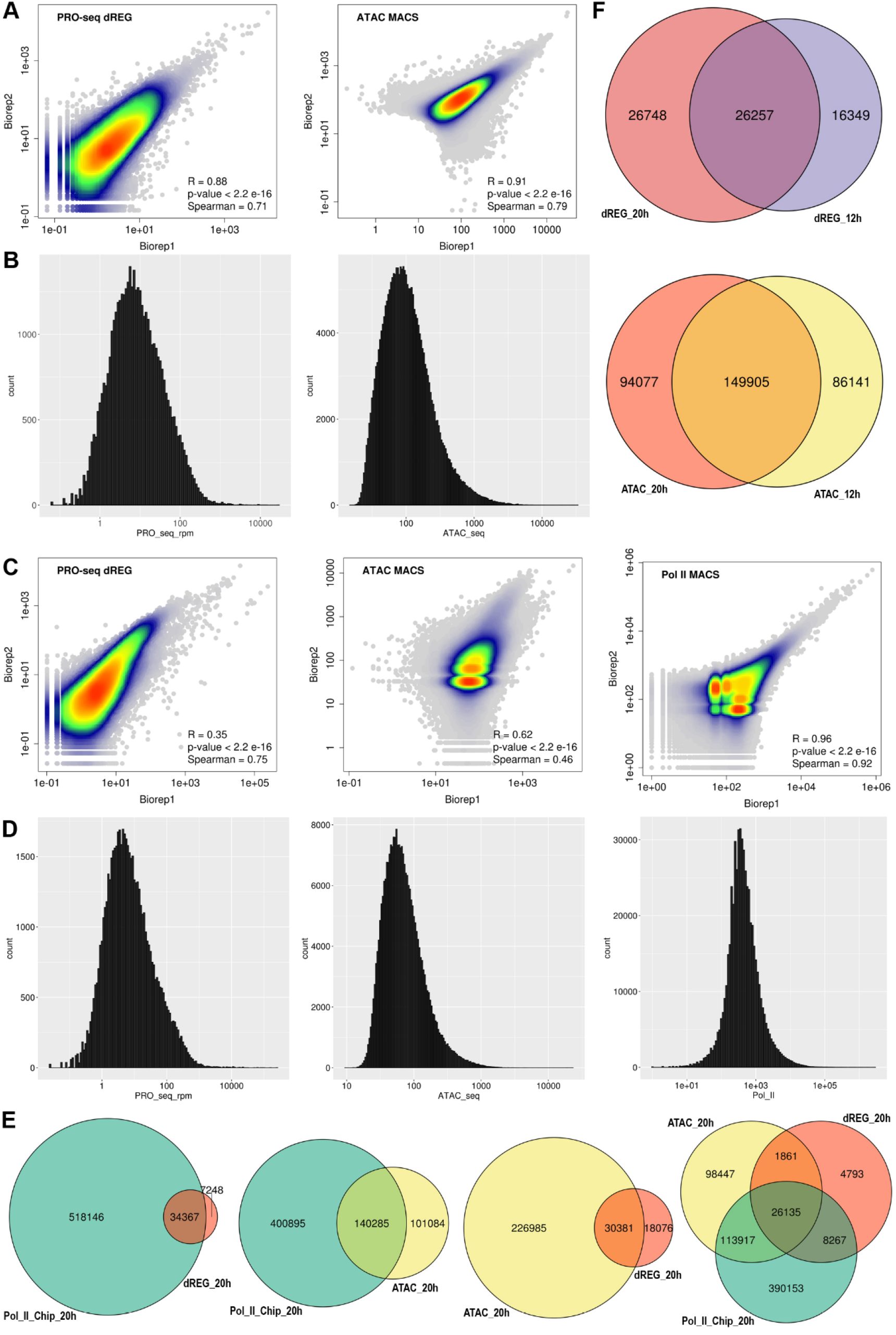
Genome-wide PRO-, ATAC- and ChIP-seq analysis. - A, Distribution of signal intensity and reproducibility estimation between distinct biological replicates for the different data sets in 12 hour embryos. Overlap of points indicated by the color gradient.
- B, Histograms of the number of reads per peak call for the different data sets in A.
- C, Distribution of signal and reproducibility in 20 hour embryos.
- D, Histograms of the number of reads per peak call for the different data sets in C.
- E, Venn diagrams of the overlap between ATAC and Pol II ChIP peak calls, and dREG predicted TREsin 20 hour embryos.
- F, Venn diagrams of the overlap of ATAC and dREG peak calls between stages.

Interestingly, there is less variability among Pol II ChIP-seq biological replicates(R = 0.96 to 0.94, with p-values < 2.2 ×10^−16^, **Fig. 2 A**). Nevertheless, the biological or technical source of the variation among the different marks could not be resolved in this study, because different embryo batches, sometimes from different seasons, were used. In addition, for the 20 hour ATAC-seq data, a distinct nuclear extraction protocol for one of the replicates (19) may have contributed to some technical variability. However, despite the higher variability of the PRO-seq and ATAC-seq 20 hour data sets, generally similar signal profiles are seen among biological replicates in both developmental stages, as illustrated at the *H2A.Z* locus (19) (**Fig. 1 B**). Similar reproducibility trends are observed at promoters and CRMs (**Fig. S1 A-D**), with much higher correlations for the subset of CRMs that are the primary target of this study (CRMs hereafter) (**Fig. S1 B and D**).

Genome-wide, most dREG peaks overlap Pol II peaks (**Fig. 2 E**), as expected. However, because much of Pol II is found in the body of transcribed genes, the majority of pol II peak calls did not overlap with dREG predictions. About 40 % of ATAC-seq peaks do not overlap Pol II peaks, and about 90 % of ATAC-seq peaks do not overlap dREG predictions, revealing that a substantial fraction of chromatin-accessible regions do not associate with RNA Pol II or transcription initiation detected using dREG, which depends on local transcription initiation profiles. Similar overlapping trends are observed in the CRMs target of this study, with a much larger fraction of ATAC peaks overlapping Pol II peaks, dREG predictions and both (**Fig. S1 E**), possibly in association with a transcriptional regulatory enrichment in the CRM data set, which is strongly biased for evolutionary sequence conservation (27).

The distinct peak numbers and particular overlaps among the three genomic assays anticipate distinct contributions and/or the requirement of combinatorial analysis for the prediction of distal TREs. Globally, about 42 % and 50 % of the 12 and 20 hour dREG peaks are stage specific, respectively (**Fig. 2F**), while 36% and 38 % of the 12 and 20 hour ATAC peaks are stage specific, respectively (**Fig. 2F**). However, for peaks overlapping CRMs (**Fig. S1 F**), the majority of ATAC and dREG peaks present in the 12 hour stage remain in the 20 hour stage, but a much larger proportion of dREG peaks than ATAC peaks are 20 hour specific, 60% versus 30 %. This reveals that during the 12 to 20 hour transition there is a general increase of transcription initiation and pause at developmental TREs while accessibility, estimated by peak calls, is more constant. This suggests that increased accessibility of developmental enhancers generally precedes transcriptional output of developmental TREs.

### Validation and evaluation of functional genomic marks for the identification of developmental TREs

We manually contrasted our functional genomic data sets with previous experimental *cis*-regulatory analyses in order to explore how they could facilitate the identification of active developmental TREs. TRE necessity for the control of endomesoderm transcription factor *SpHox11/13b* developmental expression has been characterized by deletion from BAC reporters, and TRE sufficiency by plasmid reporter constructs testing an overlapping array of genomic regions that scan the entire locus (18). ATAC-seq peaks underscore regulatory element ME in 12 and 20 hour embryos but only dREG highlights ME in 20 hour embryos (**Fig. 3**), which corresponds to the stage of higher ME reporter activity (18).

**Figure 3.**
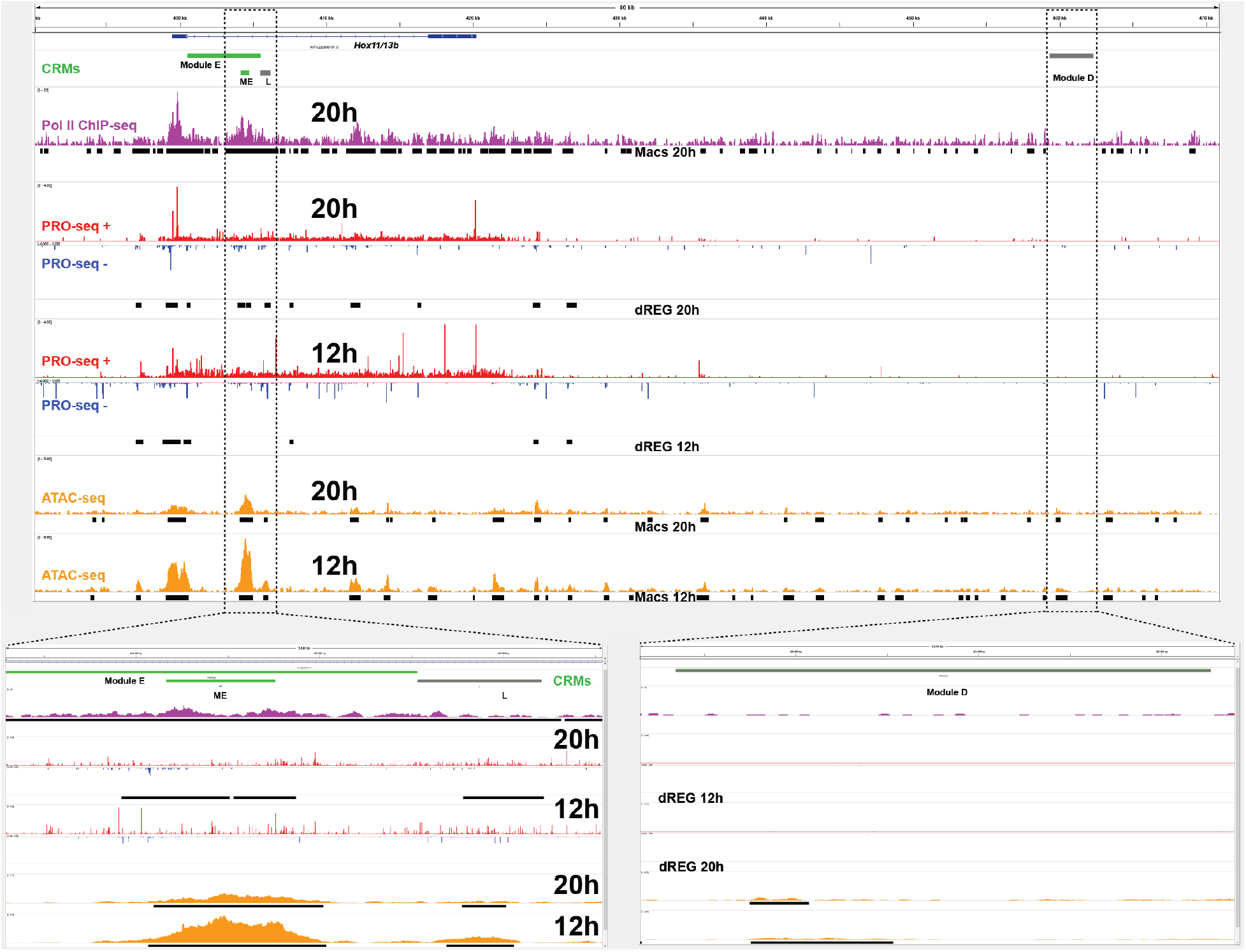
ATAC-, Pol II ChIP- and PRO-seq sea urchin embryos at the *SpHox11/13b* locus. For ATAC- and PRO-seq, the scale in reads per million at the start of each track is maintained at the same range between states and equal between plus and minus strands. The whole region was scanned for enhancer activity by overlapping 3-5 Kb reporter constructs (18), only active CRMs are indicated, in greent hose active in both stages, and in gray those inactive or with unknown activity in these stages as indicated in the text.

This pattern of 20 hour specific dREG activity follows the general ATAC and dREG stage prevalence trend in the proximity of regulatory elements, a generally constant number of ATAC peaks an increased number of dREG peaks during the later blastula stage (**Fig. S1 E**). Module ME has been demonstrated to be both necessary by deletion in BAC reporters and sufficient in plasmid reporters (18) to drive the embryonic *SpHox11/13b* expression profile (34). Like many other early embryo TREs, ME has distinct enhancer and silencer functions in different embryonic territories. In 12 and 20 hour embryos, module ME responds to spatially restricted vegetal *wnt* signaling by enhancing transcription in the endomesoderm and endoderm territories, and by silencing transcription in the ectoderm and the mesoderm, where the *wnt* pathway remains and becomes inactive, respectively (18). Therefore, the whole embryo genomic profiles derive from both enhancer and silencer activities from different territories.

DNA binding sites for transcription factor TCF are required for the *wnt* signal dependent enhancer and silencer functions of element ME. There is an increase in the transcriptional pause at the 20 hour embryo *SpHox11/13b* promoter relative to the 12 hour stage **(Fig. 3**), which could correspond with its ME transcriptional silencing in the ectoderm and mesoderm (18). Interestingly, the cofactor of TCF, groucho, implements silencing by pause in Drosophila embryos (35), which may represent an evolutionarily conserved function in sea urchins. Module D, in isolation, drives unrestricted reporter expression in 15 and 18 hour embryos that can be dominantly silenced by module ME when placed in the same reporter construct (18). Module D is inactive in 6 and 21 hour embryos, leaving uncertain its activity in 12 hour embryos. There are ATAC-seq peak calls with relatively low signal within Module D in both stages, but no dREG peaks (**Fig. 3**). Thus, module D lacks silencing functions and dREG peak calls.

Module D was not deleted in BACs and therefore its endogenous function remains uncertain (18). Finally, element L, which drives reporter activity at later stages, was undetectable with reporter constructs in 15 and 24 hour embryos (18), dREG marks regulatory element L in 20 hour embryos but not in 12 hour embryos, while ATAC detects this regulatory element in both stages. In this case, the dREG peak calls and associated pausing in module L of 20 hour embryo may correspond with its priming for subsequent activation during later embryonic and larval stages.

Similar to *SpHox11/13b* module ME, distal regulatory module Intron-D of *onecut* integrates enhancer and silencer functions that are necessary and sufficient to recapitulate the expression of this transcription factor (36). Likewise, ATAC-seq peak calls underscore Intron-D in both stages, but only 20 hour dREG peak calls highlight Intron-D, coincident with an augment of pause at the *onecut* promoter (**Fig. S2**). Thus, both in *SpHox11/13b* ME and *onecut* Intron-D PRO-seq signals may correspond to a blend of transcriptional activation and silencing functions in different embryo regions. Only ATAC-seq peaks intersect *onecut* Intron-C module, which is inactive in 20 hour embryos (**Fig. S2**). Thus, similar to module L of *SpHox11/13b,* the ATAC-seq peak call does not correspond with enhancer activity.

There are far more ATAC-seq peak calls than dREG predictions at both loci (**Fig. 3 and Fig. S2**), along with the general genomic trend (**Fig. 2. E**). Both loci where scanned by a comprehensive reporter tiling scheme to test the entire regions for enhancer activity in an unbiased manner (18,36). Remarkably, dREG TRE predictions correspond closely with regulatory elements experimentally mapped to their minimum range (**Fig. 3 and Fig. S2**). However, most dREG predictions do not match CRM enhancer reporter activity (**Fig. 3 and Fig. S2**), and even a higher proportion of ATAC peaks do not correspond with enhancer-active CRMs. The distribution of Pol II ChIP peaks is even broader, particularly at introns, which are prone to contain TREs **(Fig. 3 and Fig. S2**), and, therefore, Pol II ChIP signal seems poorly suited for TRE predictions on its own. Manual analysis of other experimentally characterized transcriptional regulatory elements (37–39) generally confirms the trends outlined above (**Fig. S2**). Additional regions of the genome can be analyzed with the data sets deposited at the NCBI Gene Expression Omnibus (40). In summary, our manual analysis suggest that accessibility may have a looser correspondence with enhancer activity. In addition, although increased pause does not necessarily correspond with silencing, as it may be associated with increased release and elongation, the dual report of transcriptional elongation and pause by PRO-seq may correspond to enhancer and silencer activities differentially deployed in space, which is very common among early embryo regional specification TREs (4).

### Prediction of enhancer activity from chromatin accessibility and Pol II

We decided to systematically test if machine learning models using chromatin accessibility, Pol II distribution and transcription initiation could predict the previously quantified enhancer activity of 389 CRMs primarily selected for their evolutionarily sequence conservation (27). We first tested a subset of reporters quantified at high temporal resolution using nanoString technology (26), but the very small number of inactive reporters was unsuitable for model training (results not shown). We therefore settled for a previous data set that measured reporter enhancer activity by qPCR and included 12 and 24 hour time points (27). Although this generated a mismatch with the 20 hour stage examined in our genomic data, no major regulatory transitions have been identified for the majority of the genes involved during this 4 hour period (41). The average size of the 389 CRMs (2,839 bp) is about one order of magnitude larger than the average of ATAC peaks (316 bp) Pol II peaks (338 bp), and dREG TRE predictions (373 bp) **(Fig. 4 A**). Thus, in order to reduce confounding inputs to the CRMS that may not relate to TRE function, such as background transcription at introns, or background accessibility along the CRM span, the ATAC-, Pol II ChIP- and PRO-seq signals were computed at CRM regions overlapping peak calls and dREG predictions. CRMs were defined as active if they drove reporter expression twice above the basal promoter (**Fig. 4 B and C**). CRM activity cannot be explained by CRM size because there is no significant size difference between active and inactive enhancers (**Fig. 4 A**, inset, Wilconox p-value = 0.11). However, active CRMs in 12 and 24 hour embryos have significatively higher PRO-, ATAC-, and Pol II ChIP-seq signals (**Fig. 4 D**, p-values < 1.8 e-06).

**Figure. 4.**
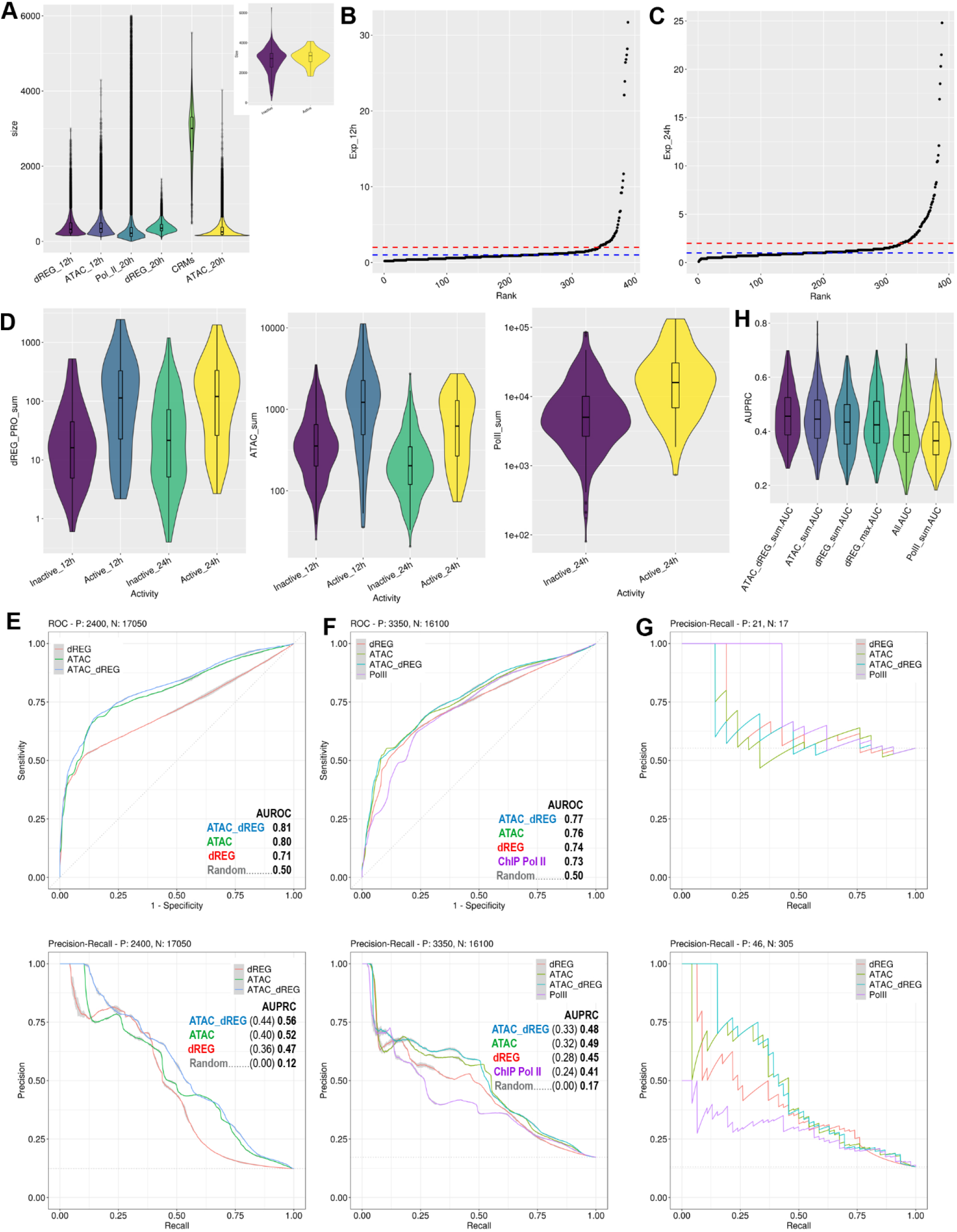
Modeling of CRM reporter activity from of ATAC-, Pol II ChIP- and PRO-seq. - A, Violin/box-plot of the ATAC, Pol II ChIP peak call and dREG TRE prediction sizes, and the 389 CRMs. The inset plots the size distributions of active and inactive CRMs, which is not significatively different.
- B and C, ranked CRM expression plot in 12 and 24 hour embryos, respectively. The blue line at 1 marks the CRM expression level when it equals that of the basal-promoter reporter. The red line by the curve “elbow” marks the 2 fold above control chosen as the expression threshold.
- D, Violin/box-plots of PRO-, ATAC-, and Pol II ChIP-seq significatively different signals between active and inactive CRMs in 12 and 20 hour embryos.
- E, top, 12 hour embryo Receiver Operating Characteristics (ROC) and, bottom, 3 iteration average Precision-Recall Curves (PRC) of the logistic regression models trained and tested by 5 fold cross-validation repeated 50 times. Abutting each line, 95% confidence interval bands estimated after the 3 model iterations are shown in grey, see methods for interpretation. Area Under the ROC (AUROC) and AUPRC as indicated for each model. Dottedlines mark random guess prediction performance, a mid-diagonal for ROC and a horizontal line at the fraction of active CRMs for PRC. The absolute AUPRC indicated in bold and the difference with random guess in parenthesis.
- F, ROCs and PRCs in 20 hour embryos.
- G, top, PRCs evaluating the enhancer activity predictions for the CRM promoter- overlapping data set of models trained with the entire 20 hour CRM data set. Bottom, model predictions for the complementary, non-promoter overlapping data set.
- H, Violin/box-plot of the AUPRC after cross-validation with different predictors, as indicated; All, includes the sum and max of the 3 genomic profiles allowing second order interactions among predictors; dREG-max, signifies the sum of the maximum values at dREG peaks.

Logistic regression classifiers trained and tested by 5 fold cross-validation repeated 50 times resulted in predictions significatively above random guess in both embryo stages (**Fig. 4 E and F**). The performance of models using ATAC, dREG (PRO-seq reads at TRE dREG predictions), their combination (ATAC + dREG), and Pol II ChIP was slightly higher for ATAC + dREG models when evaluated by the Area Under the Receiver Operating Characteristic (AUROC) plot (**Fig. 4 E and F**), which graphs the relation between true and false positive rates at different model prediction thresholds. However, the CRM expression data set is highly unbalanced, with about 10 times more CRMs reporting inactive than active enhancer activity **(Fig. 4 B and C**), and, in these cases, Precision-Recall Curves (PRCs), which plot precision values along the range of true positive rates, provide a better discrimination metric for classifier evaluation (42).

When AUPRCs are used for model evaluation, more distinct model performances are obtained, particularly for the 20 hour data sets (**Fig. 4 F, bottom**). Individually, all assays perform much better than chance in both stages (**Fig. 4 E and F**). The combination of ATAC and dREG predictors may slightly improve performance at some recall values (**Fig. 4 E and F, bottom**), and Pol II ChIP signal does not facilitate better enhancer activity predictions alone (**Fig. 4 F**) or in combination with other data sets (results not shown). Incorporation of other parameters such as peak summit value, did not improve any predictive models, as illustrated for dREG (**Fig. 4 H**). As expected, lower model performance was also obtained when the functional genomic data were computed along the entire CRM instead of restricting the signal input to peak call windows (not shown). Optimization of other machine learning methods, such as random forest and support vector machine, did not improve classifier performance over logistic regression (not shown), likely reflecting the small size of the available data. In short, total ATAC-seq and PRO-seq signals at dREG peaks are the best predictors of active enhancer activity among the profiles tested in this study.

Interestingly, about half of the CRMs that overlap promoters are active in the reporter assays, indicating a high degree of enhancer activity from promoter-adjacent DNA in sea urchin embryos. Nearly all these promoter-overlapping CRMs were previously shown to be active in both orientations (27), demonstrating bona fide enhancer activity. The sizes of promoter-overlapping CRMs (**Fig. 4 A**) suffice to include both distal and proximal TREs, including promoters. ATAC and dREG models trained with the entire data set (**Fig. 4 F**) underperformed relative to Pol II ChIP based models in the prediction of enhancer activity of the promoter-overlapping CRM subset (**Fig. 4 G, top**). In the complementary analysis, the Pol II ChIP model trained with the entire set further underperformed relative to ATAC and dREG models in the prediction of CRMs not overlapping promoters, while ATAC and dREG maintained performance similar to predictions with the entire set (**Fig. 4 G, bottom**). The exclusion of the 41 promoter-overlapping CRMs from the training and testing data set decreased the prediction performance of all models in both stages(**Fig. S3 A and B**). Overall performance was broadly similar between ATAC and dREG models trained and tested with the promoter overlapping or non-overlapping (**Fig. S3 A and C**). In contrast, ATAC and dREG models trained with CRM-overlapping promoters failed to predict the activity of their hold out set and where outperformed by Pol II ChIP models in the 20 hour data set (**Fig. S3 D**). All of the above, suggests that distinct functional marks associate with the enhancer activity of promoter proximal and distal TREs, and that the enhancer activity predictive power of ATAC-seq and PRO-seq for promoter-proximal CRMs dramatically devalues during the 12 to 20 hour transition.

The larger proportion of positive enhancers among CRMs that overlap promoters relative to CRMs not overlapping promoters, ∼ 50 % vs. ∼13% in 20 hour embryos, is not surprising given the bias for regulatory genes active during development of this data set (27) combined with the general trend of enhancers to be near their promoter targets(43). The enhancer activity of promoters has precedents (44,45) and it is perhaps not surprising for evolutionary reasons (2).

We tested if the functional genomic datasets could predict the levels of reporter enhancer activity of CRMs. In all cases, better linear regression model prediction was obtained with non-promoter overlapping CRM sets. The best performing model included the ATAC-seq plus the ATAC: dREG interaction, which explained about one third of the expression variation (average R^2^ = 0.29) in 20 hour embryos (**Fig. 5**). ATAC-seq was a better predictor or enhancer activity in 20 hour embryos relative to 12 hour embryos (R^2^ = 0.26 versus R^2^ = 0.17, p-value < 2.2 e-16), and outperformed dREG in 20 hour embryos (R^2^ = 0.17, p-value = 7.4 e-16) (**Fig. S4**). The difference in dREG model performances between stages or with ATAC-seq models in 12 hour embryos was not significant due to the small size of the dataset. Nevertheless, the relative enhancer predictive power of ATAC-seq and PRO-seq is stage-dependent. The predicted value of most CRMs generally varies with the training set, as expected. However, there is a group of CRMs that are consistently and erroneously predicted as barely active (**Fig. S4**) due to their low signals in all assays (not shown). Highly active reporter constructs with low predictor signals may result from not uncommon miss-regulation outside the endogenous genomic context (4), which may cause ectopic expression, such as the one observed for *SpHox11/13b* module D (18), or it may reflect mismatches between the time points, especially at the 20 hour stage.

**Figure 5.**
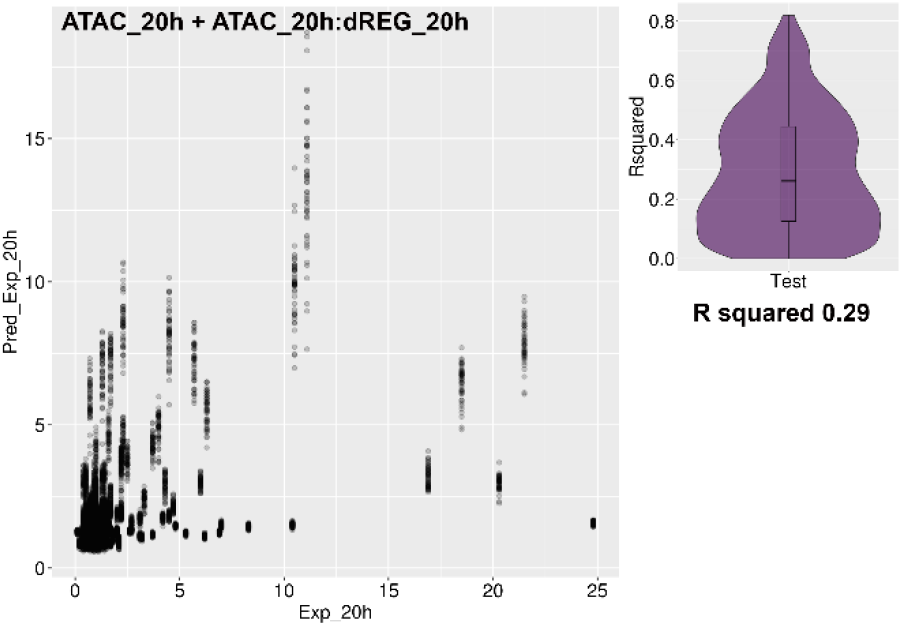
Quantitative prediction of enhancer activity from PRO-seq data. Plot of the hold-out predicted against the actual reporter expression of linear regression models using ATAC and PRO-seq signal at dREG predictions tested by five-fold cross-validation. Violin/Box-plot of R^2^ values, with the average indicated underneath.

## Discussion

Machine learning models identify ATAC- and PRO-seq as efficient predictors of the developmental enhancer activity of genomic regions previously validated by their reporter driven expression in sea urchin embryos (**Fig. 4 E and F**). Further prediction improvements are expected after addressing some limitations of our experimental setup. The bulk functional genomic profiles of whole embryos represent a blend of several transcriptional states present in different territories. The unavoidable bias against regulatory elements only active in a few cells should nevertheless correspond with similarly biased CRM reporter expression levels. The mismatch between the size of TRE peak calls and CRMs tested is less than ideal. In most high-throughput reporter assays (11), the regulatory regions tested are usually smaller than the few hundred base pairs of common TREs, which represents one of the several limitations of enhancer activity evaluation by reporter constructs (11). In contrast, the genomic regions tested in our data set are large (**Fig. 4 A**) and often contain several ATAC and dREG peak calls, whose signals would possibly better match enhancer activity if individually tested. In addition, enhancer activity is tested with a heterologous promoter, allowing for the mismatch between functional genomic assays and reporter activity due to enhancer-promoter specificity (46).

Despite the tight match between dREG TRE predictions and CRMs experimentally narrowed down to the smallest functional regulatory elements in a generally unbiased manner (**Fig. 3**), our results reveal that PRO-seq has similar predictive power as ATAC-seq. Perhaps this results from the dual report of transcription and pause by PRO-seq. In addition, enhancer and silencer activities in different territories are common for developmental regulatory elements (4,47), as previously discussed in the context of *SpHox11/13b* and *onecut* (**Fig. 3 and Fig. S2**). Alternatively, the small set of positive enhancers measured by reporter assays could result in having insufficient statistical power. Nevertheless, despite all these caveats, ATAC-seq and PRO-seq alone suffice to explain between one quarter and one fifth of the reporter enhancer activity in 20 hour embryos (**Fig. 5 and Fig. S4**). It is reasonable to expect even better performance in single cell assays exclusively testing the genomic regions highlighted by ATAC- and PRO-seq profiles.

Our results confirm and extend reports of distinct enhancer prediction performance for promoter-proximal regulatory elements previously obtained with a distinct set of functional genomic profiles (14). More interestingly, ATAC and PRO-seq profiles have similar predictive power for promoter-overlapping and distal CRMs in 12 hour embryos, but become irrelevant for the prediction of promoter-overlapping CRMs in 20 hour embryos, when only Pol II has predictive value **(Fig. 4 G and Fig. S3 D)**. This suggests that distinct transcriptional regulatory mechanisms may prevail at TREs used at different levels of the gene regulatory hierarchy that is sequentially deployed during development, or that regulatory genes are at distinct stages in the sequence of events leading to their transcriptional activation.

Major regulatory changes impact the 37 regulatory genes subject of this study during the 12 to 20 hour transition, as summarized by the topological models of the BioTapestry Interactive Network Viewer (41). In short, the activation of the upstream regulatory genes that determine the main territorial subdivisions is underway in the 12 hour blastula, while the transcriptional states determining these subdivisions are well stablished in the 20 hour blastula, including the activation of terminal differentiation gene batteries at the periphery of the transcriptional network (15–17,41,48). In general, more PRO-seq than ATAC-seq associated changes are observed during the 12 to 24 hour transition (**Fig. S1 F**), in agreement with punctual observations (**Fig. 3 and Fig. S2**). This would follow the general sequence of events in the transcriptional cycle (49), with enhancer accessibility preceding transcriptional initiation, followed by pause and release, all of which are target of regulation by sequence specific transcription factors. The early territorial subdivisions are mediated both by transcriptional enhancer and silencer functions, and the relation of functional genomic profiles to experimental characterizations (**Fig. 3 and Fig. S2**) suggest that pausing may not only provide a venue for coordinated and prompt transcriptional activation during development (50), but also anticipate permanent silencing in some territories.

## Conclusions

In summary, ATAC- and PRO-seq are efficient predictors of reporter enhancer activity of distal CRMs in sea urchin embryos, while the prediction of promoter-overlapping CRMs is stage-dependent. In late blastula embryos, Pol II enrichment is the best predictor of promoter-proximal CRM enhancer activity. There is a net increase in dREG TRE predictions during later embryonic stages, while accessibility peaks remain relatively constant. In combination, this suggests that the sequence of regulatory events leading to developmental TRE enhancer activity has different relevance at different GRN levels or developmental stages. Our work facilitates ongoing developmental gene regulatory studies by mapping genome-wide candidate TREs, identifies PRO-seq and ATAC-seq as candidate factor-independent methods that predict developmental enhancer activity in whole embryos, and outlines the stage-dependency and predictive value of distinct functional genomic profiles associated with proximal and distal regulatory elements.

## Methods

### Preparation of nuclear extracts and sequencing libraries

Sea urchin embryos were reared to different stages as previously described (19). Nuclei for ATAC-seq and PRO-seq were prepared using a modified version of a density gradient method (51) as follows. Sea urchin embryos were centrifuged at 500 g for 3 minutes at 0 °C, the pellet was resuspended in 10 volumes of ice cold lysis buffer consisting of 20 mM EDTA, 2% polyethylene glycol, and 4 mg/ml of Protease Inhibitor Tablets (Thermo Scientific™ Pierce™ #A32965), added just before use, in 0.1 X PBS (PBS is 0.137 mM NaCl, 2.7 mM Cl, 10 mM Na_2_HPO_4_ and 18 mM KH_2_PO_4_), and incubated on ice for 5 minutes. Dissociated cells were further disrupted with 50 or more strokes in a fine dounce homogenizer. Density gradient nuclear wash and floating layers were prepared by diluting iodixanol 60 % (OptiPrep™) in 1 X PBS to 20 % and 40%, respectively. About 5 ml of nuclear lysate was deposited on top of 10 ml of nuclear wash and nuclei were collected over 200 µl of floating layer after centrifugation for 30 minutes at 2 °C and 3,000 g in a swing bucket rotor. Nuclei aliquots were flash-frozen in liquid nitrogen. For the 20 hour stage, nuclei of one of the two ATAC-seq biological replicates was prepared as previously described (19). For ChIP-seq, fixation was performed by resuspending embryo pellets in crosslinking solution (1 mM EDTA, 0.5 mM EGTA, 100 mM NaCl, 1.8 % formaldehyde, 50 mM HEPES, pH 8.0) for 15 minutes at 22 °C, followed by gravity settling and subsequent resuspension in stop solution (125 mM glycine, 0.1% Triton X-100 in PBS), 500 g centrifugation, and two washes with PBT (0.1% Triton X-100 in PBS). The embryos were transferred to 25 ml of ice cold homogenization buffer (15 mM Tris-HCl pH 7.4, 0.34 M sucrose, 15 mM NaCl, 60 mM KCl, 0.2 mM EDTA, 0.2 mM EGTA, with 4 mg/ml protease inhibitors) and incubated on ice for 5 minutes. Embryos were first dounced 20 times with pestle type A (loose), followed by 10 times with pestle type B (tight). Nuclei were then filtered through a 20 μM filter and pelleted at 3,500 g for 5 minutes at 4 °C. The nuclei were resuspended in 7.5 ml of PBTB (5% BSA in PBT buffer, with proteinase inhibitors). Propidium iodine stained nuclei were quantified using a hemocytometer and a fluorescence microscope.

ATAC-seq library preparation and Illumina sequencing followed similar procedures to those previously described (28). The ENCOCE-DCC atac-seq-pipeline (52) was used for mapping the raw reads to the *Strongylocentrotus purpuratus* genome version 3.1 using default settings, except for the MACS2 peak call p threshold, which was set to 0.05. Two biological and one technical replicate libraries were paired or single end sequenced. A total of 16,005,927 20-hour and 73,272,494 12-hour embryo reads mapped to the genome after deduplication and mitochondrial chromosome filtration. About 44 % of the reads locate in peak calls, see quality control summary for detailed reproducibility and sequence quality metrics (Supplementary quality control files).

PROseq libraries were elaborated following previously stablished protocols (53), single or paired end sequenced in Illumina platforms, and mapped to the *Strongylocentrotus purpuratus* genome version 3.1 using proseq2.0 pipeline (54). Regulatory elements were predicted with the vector machine learning tool dREG (10). Two biological and one technical replicate libraries were prepared and paired or single end sequenced, providing a total of 56,781,051 20-hour and 28,430,031 12-hour mapped reads, excluding ribosomal RNAs.

For ChIP-seq library preparation, 50 to 100 million nuclei were spun at 4,000 g for 5 minutes at 4°C. Nuclei were resuspended in 1 ml FA buffer (50 mM HEPES/KPH pH 7.5, 1 mM EDTA, 1% Triton X-100, 0.1% sodium deoxycholate, 150 mM NaCl, 0.1% sarkosyl, and protease and phosphatase inhibitors). The resuspended nuclei were then sonicated at 4°C for 15 minutes on high, cycle of 30 seconds on and 30 seconds off, to obtain an extract with fragmented chromatin. Extracts were brought up to 440 μL with FA buffer with protease and phosphatase inhibitors. Between 1 to 2 mg of extract and 4 μg of antibody were used per ChIP. Prior to the addition of antibody, 5% of the extract was taken for input. Mouse monoclonal antibody against RNA polymerase II CTD-repeat YSPTSPS (8WG16; abcam ab817mod) was used for ChIP. The mixture was incubated rotating at 4°C overnight. 40 μL of protein G sepharose bead slurry (GE Healthcare) per ChIP sample was washed three times with 1 mL FA buffer, added 40 μL bead slurry to each ChIP sample and rotated at 4°C for 2 hours. Meanwhile, 200 μL ChIP Elution Buffer (1% SDS, 250 mM NaCl, 10 mM Tris pH 8.0, 1 mM EDTA) and 2 μL 10 mg/μL RNase A were added to inputs and incubated at room temperature. Beads were washed at room temperature by adding 1 mL of each of the following buffers and collecting beads by spinning for 1 minute at 2500 g: two times FA buffer for 5 minutes, one time FA-1 M NaCl for 5 minutes, one time FA-500 mM NaCl for 10 minutes, one time TEL buffer (0.25 M LiCl, 1% NP-40, 1% sodium deoxycholate, 1 mM EDTA, 10 mM Tris-HCl, pH 8.0) for 10 minutes, two times TE for 5 minutes. Proteinase K was added to both inputs and ChIPs and incubated in a 50°C heat block for an hour. Inputs and ChIPs were allowed to reverse crosslink overnight in a 65°C water bath. DNA was ligated to Illumina or homemade multiplexed adapters and amplified by PCR. Using a thin 1.5% agarose gel, DNA fragments between 300 and 600 bp were purified using the Qiagen Gel Extraction kit. Qubit flourometer was used to measure DNA concentration. Single-end sequencing was performed for the ChIP-seq and input DNA at the New York University Center for Genomics and Systems Biology high-throughput sequencing facility. We combined replicates and aligned 50 bp single end reads to the *S. purpuratus* genome version 3.1 linear scaffolds using Bowtie 2 version 2.2.3 (55) with default parameters. A total of 13,762,893 ChIP and 26,476,643 input mapped reads were obtained. Mapped reads from ChIP and input were used to call peaks and coverage per base using MACS version 1.4.2(56) with default parameters.

### Computational analysis of PRO-, ChIP-, and ATAC-seq peak calls and machine learning, enhancer activity prediction

Signal at PRO-, ChIP-, and ATAC-seq peak calls was quantified using the R package bigWig (57). ATAC and PRO-seq reads of 12 and 20 hour embryos where normalized to reads per million per base. For ATAC-seq peak calls, any overlapping peaks were merged prior to analysis. Density plots used R lift posted in StackOverflow (58). Overlaps among PRO-, ChIP-, and ATAC-seq peak calls were analyzed and illustrated with ChIPpeakAnno (59) using default parameters. Promoters are defined as the 200 bp region centered at the 5’ end of transcript based gene models, which are a better approximations than GLEAN models (33). For all data sets, reads from different replicates were merged into single bigwig files and reads computed at peak calls and dREG predictions using the bigWig interface (57).

Using bedtools (60), the intersections between dREG predictions and CRMs were merged, to correct for CRM overlaps, and then extended 50bp, to compute the pause associated PRO-seq reads oftentimes extending beyond the raw dREG prediction. The total number of 3’ end reads in the plus and minus strand and the summit for each TRE prediction was estimated with the sum and max parameters of the bigWig query function. Similar analysis was performed for the reads per base for the intersection with ATAC- and Pol II ChIP peaks, without the 50 bp extension. Graphics were elaborated with ggplot and tidyverse (61).

The package caret was used for the optimization, test and evaluation of classification and regression models (62). Logistic classification and linear regression models were fitted and tested by 5 fold cross-validation with stratified sampling repeated 50 times. The package precrec (63) was used to generate ROCs and PRCs. The 95 % confidence interval abutting in gray the average PR and ROC curves (**Fig. 4 and Fig. S3**) was calculated after 3 iterations of the 5 fold cross-validation repeated 50 times. Given the extensive resampling involved, this confidence interval provides an estimation of the reliability of the average curve rather than a significance boundary for model comparisons.

## List of abbreviations

ATAC-seq: Assay for Transposase-Accessible Chromatin using sequencing
AUPRC: Area under the Precision-Recall Curve
AUROC: Area under the Receiver Operating Characteristic Curve
BAC: Bacterial Artificial Chromosome
ChIP-seq: Chromatin Immunoprecipitation followed by sequencing
CRM: *cis-*regulatory module
dREG: Discriminative regulatory-element detection from GRO-seq
GRN: Gene regulatory network
MACS: Model-based Analysis for ChIP-seq
Pol II: RNA polymerase II
PRC: Precision-Recall Curve
PRO-seq: Precision nuclear run-on sequencing
ROC: Receiver Operating Characteristic Curve
TRE: Transcriptional regulatory element
TSS: Transcription start site

## Ethics approval, accordance and consent to participate

Not Applicable

## Consent for publication

All authors read and approved the final manuscript.

## Availability of Data and Materials

The datasets generated and analyzed during the current study are available at NCBI GEO under accession number GSE160463 (40).

https://www.ncbi.nlm.nih.gov/geo/query/acc.cgi?acc=GSE160463

## Authors’ contributions

César Arenas-Mena, Sofija Miljovska, Edward J. Rice, Justin Gurges, Tanvi Shashikant, Sevinç Ercan, Charles G. Danko

CA-M designed and optimized the nuclear extraction protocols, performed the density gradient nuclear preps, various PRO-seq experiments, the ATAC-seq replicates of 12 hour embryos, the genomic mapping of ATAC and PRO-seq reads and subsequent processing, the manual curation of genomic profiles, contributed to the design and executed the machine learning analysis and wrote the manuscript and figures. SM performed the Pol II ChIP-seq experiment and the ATAC-seq replicates in 20 hour embryos. EJR performed the first PRO-seq replicate in 20 hour embryos and assisted in the elaboration of 12 and 20 hour embryo replicates. JG participated in the optimization of early nuclear extraction protocols and obtained the nuclear preps for the first round of 20 hour ATAC-seq replicates and the ChIP-seq nuclear preps. TS performed the first round of ATAC-seq genomic mapping and ChIP-seq and ATAC-seq peak call analysis. SE designed and supervised the ATAC-seq experiments in 20 hour embryos and Pol II ChIP-seq experiments. CGD supervised and analyzed the first PRO-seq replicate in 20 hour embryos, contributed to the conceptual design of the machine learning model approach, supervised all computational analysis and statistical methods, and supervised the illustration and writing of the manuscript. All authors read and approved the final manuscript.

## Acknowledgements

We would like to thank Jongmin Nam for providing the dataset sassociated with his CRM quantitative analysis, Charles A. Ettensohn for facilitating the contribution of Tanvi Shashikant while working in his lab, the Bioinformatics High Performance Computing personnel at Cornell University for the technical assistance received, all members of the Danko lab for their help and Professor Zhong Wang in particular for his assistance during the early phases of the computational analysis.

---

**Supplementary Figure 1.**
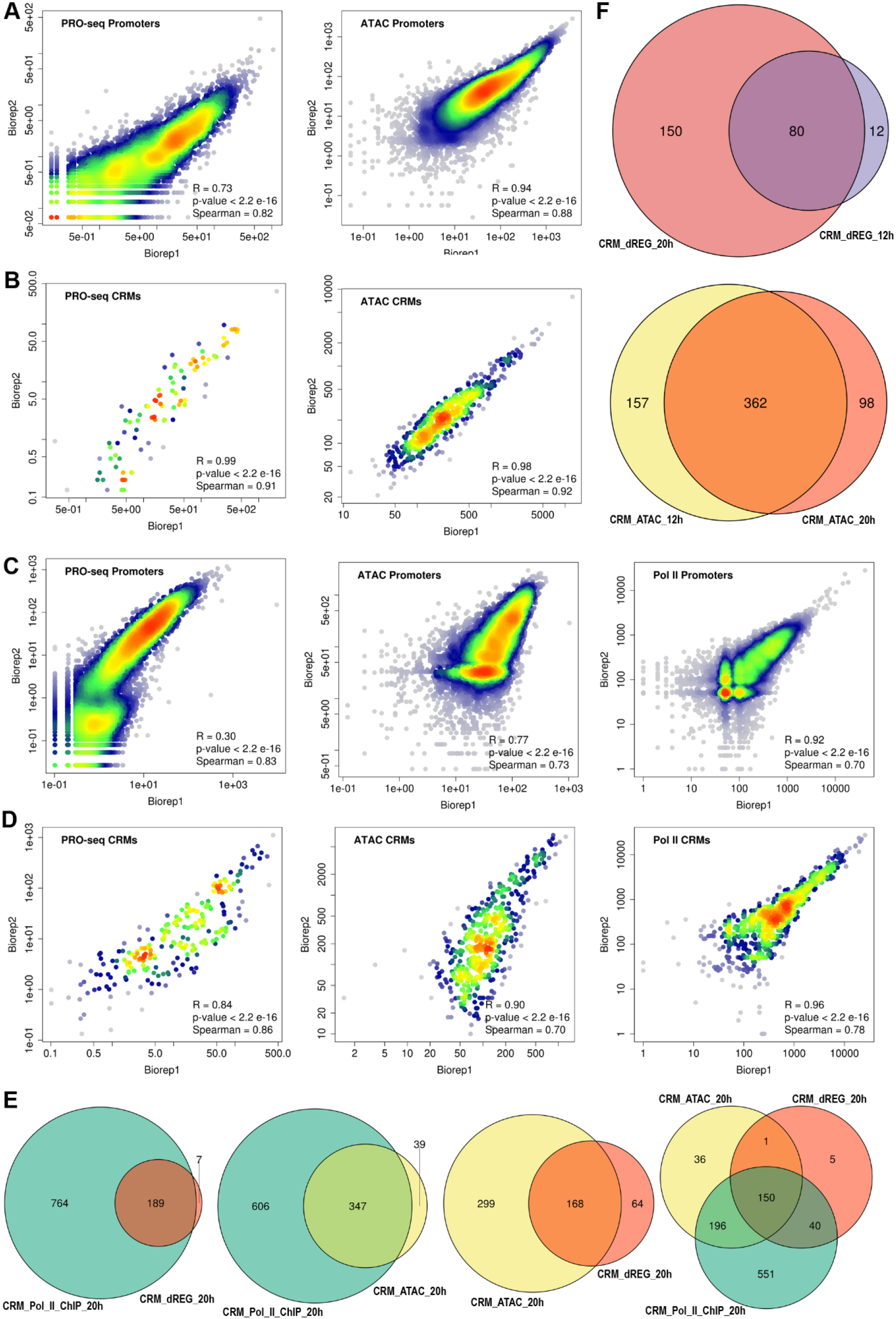

**Supplementary Figure 1.**
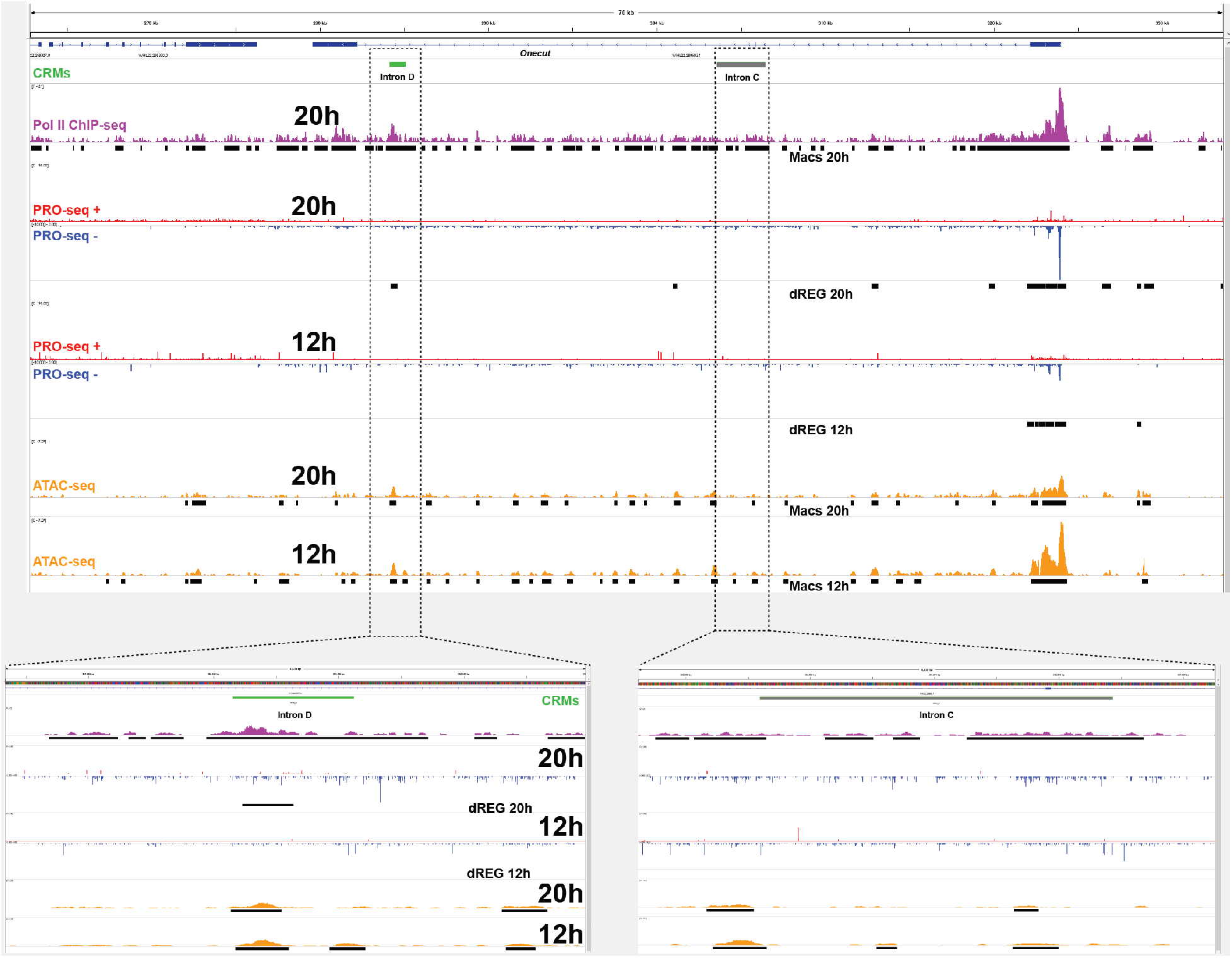
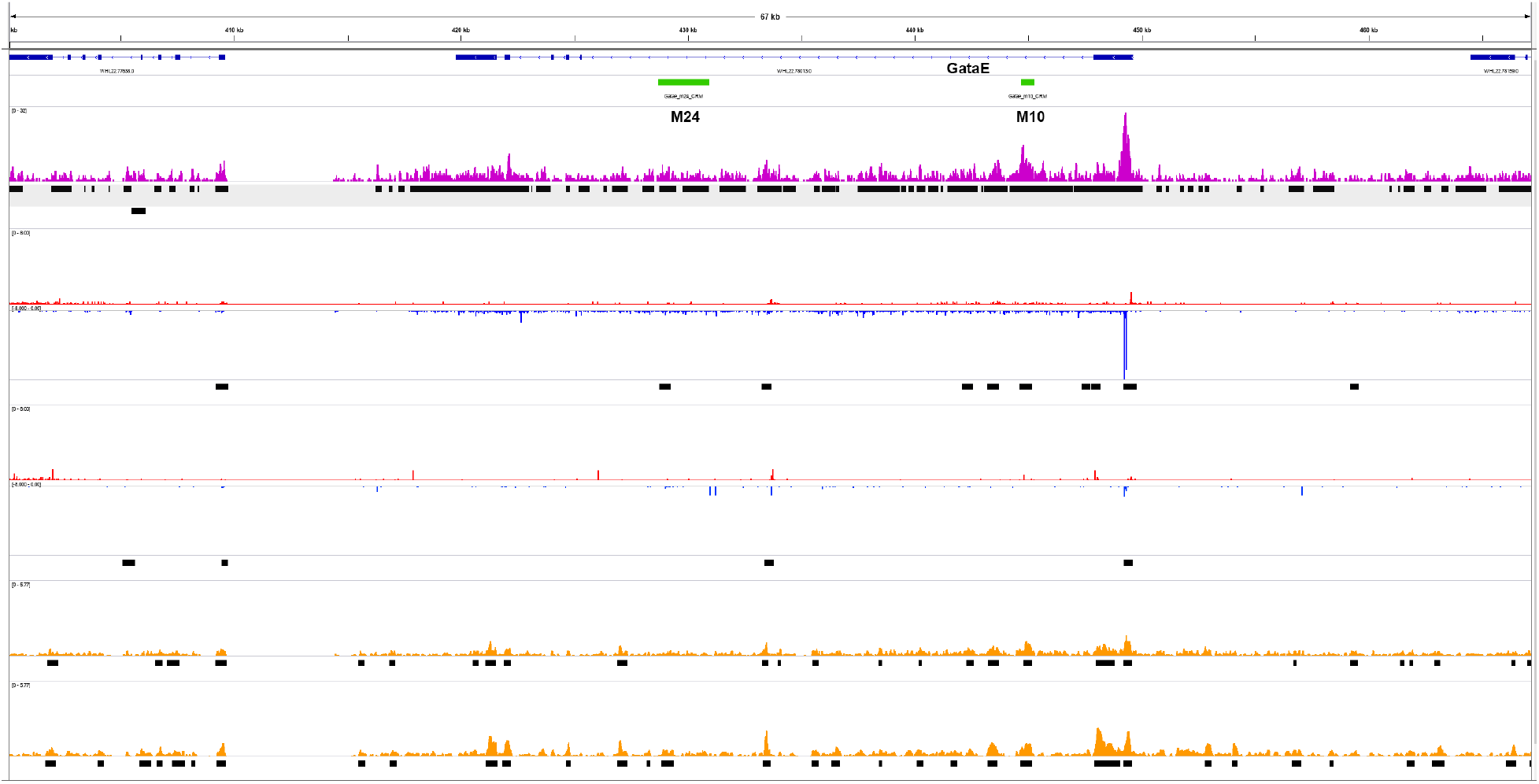
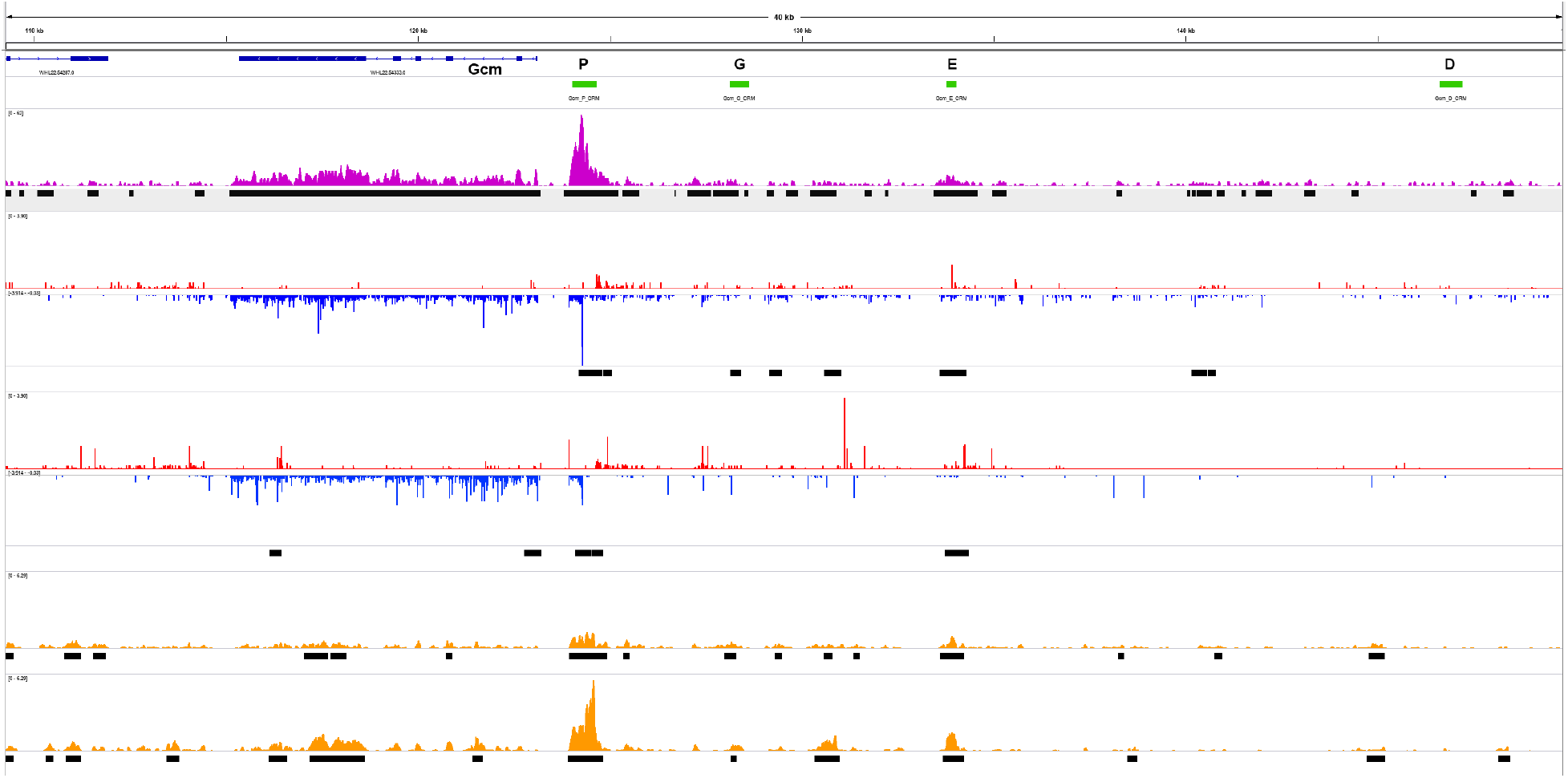

**Supplementary Figure 3.**
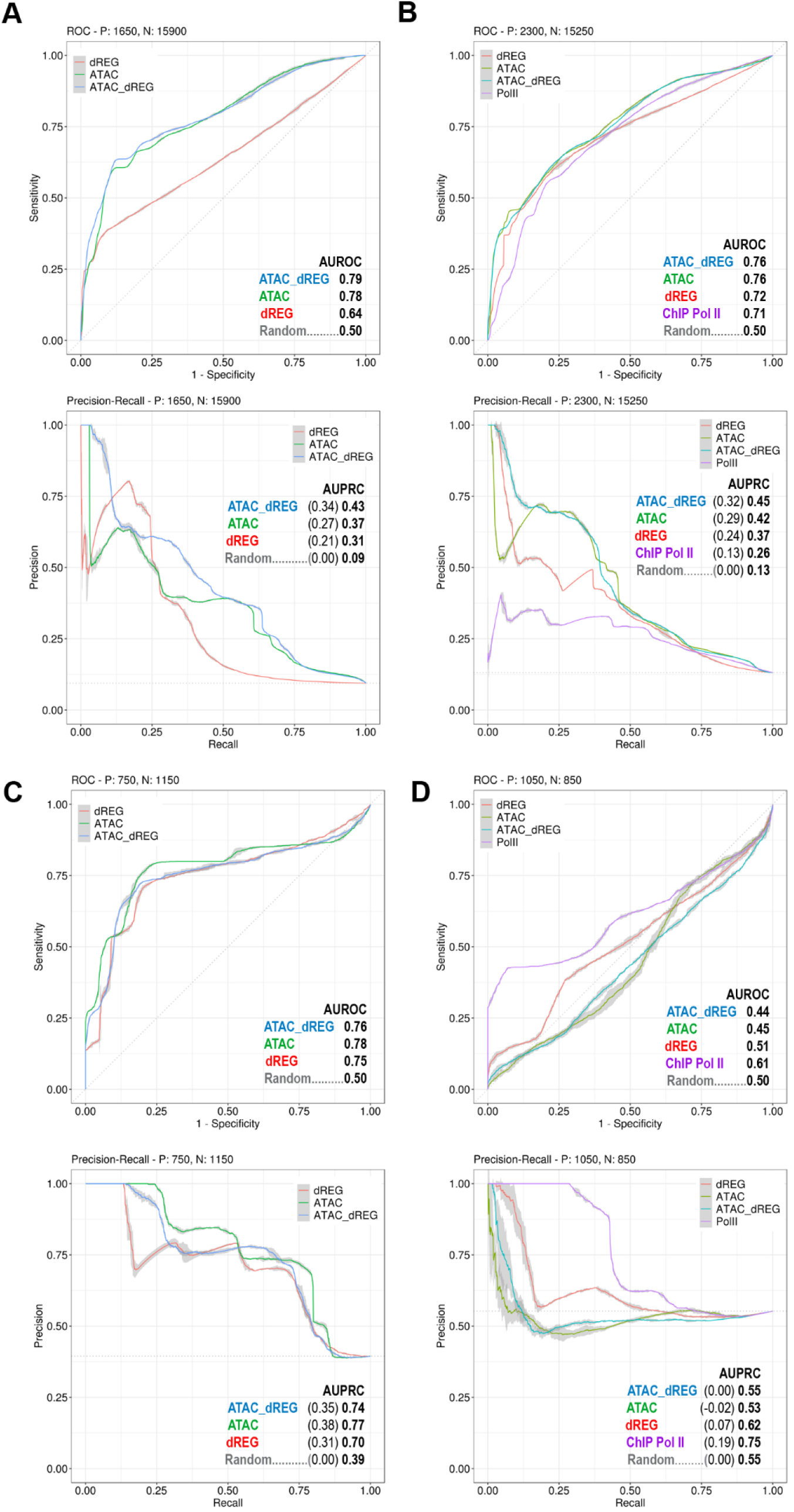

**Supplementary Figure 4.**
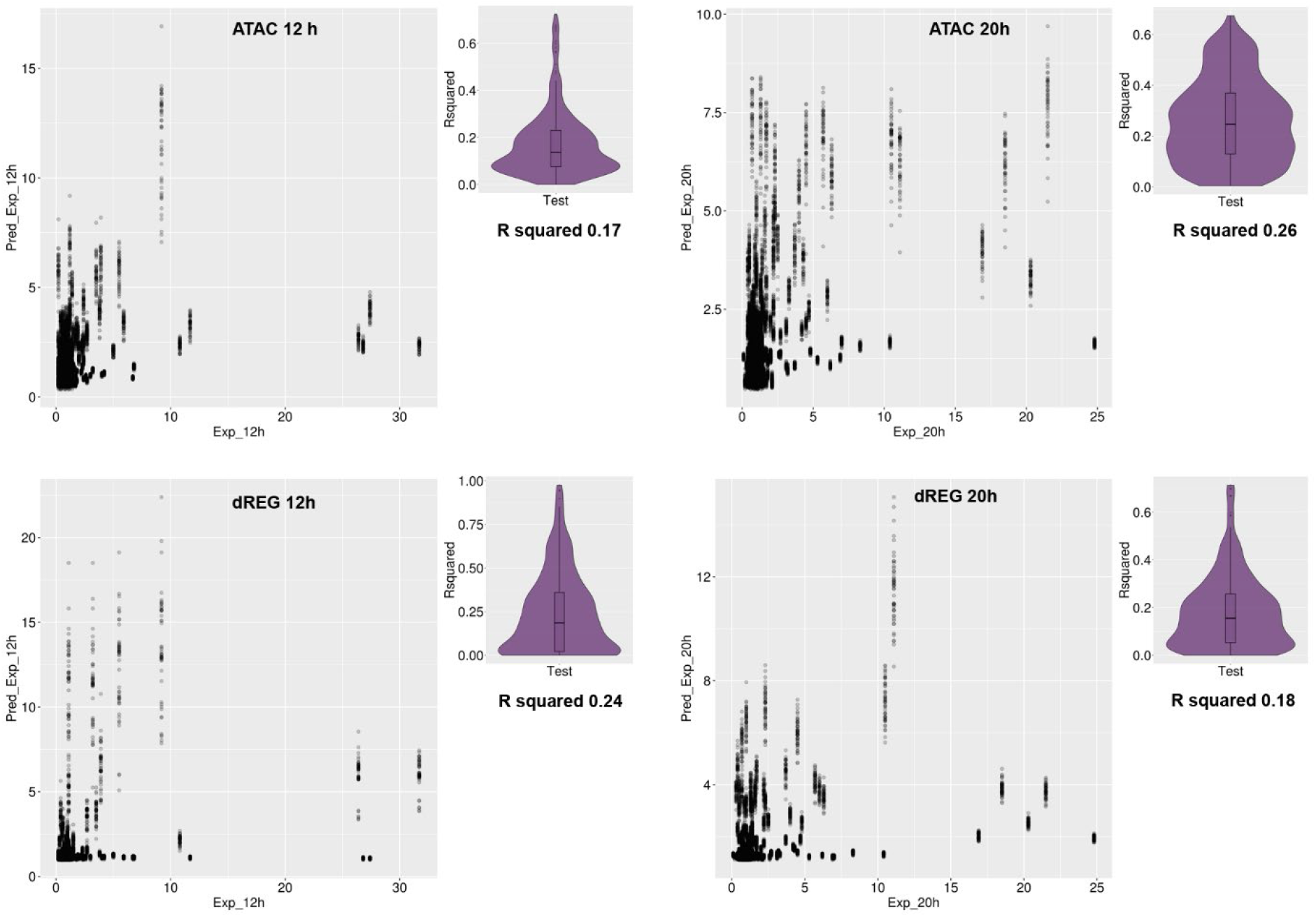

## QC Report

|  | general |
| --- | --- |
| Report generated at | 2020-10-23 01:02:22 |
| Title | ATAC_12h_3.1 |
| Description | PE 12h |
| Pipeline version | v1.7.0 |
| Pipeline type | atac |
| Genome | Sp3.1 |
| Aligner | bowtie2 |
| Sequencing endedness | {'rep1': {'paired_end': True}, 'rep2': {'paired_end': True}} |
| Peak caller | macs2 |

## Alignment quality metrics

### SAMstat (raw unfiltered BAM)

Alignment quality metrics SAMstat (raw unfiltered BAM)
|  | rep1 | rep2 |
| --- | --- | --- |
| Total Reads | 161870122 | 141132800 |
| Total Reads (QC-failed) | 0 | 0 |
| Duplicate Reads | 0 | 0 |
| Duplicate Reads (QC-failed) | 0 | 0 |
| Mapped Reads | 99587727 | 111643482 |
| Mapped Reads (QC-failed) | 0 | 0 |
| % Mapped Reads | 61.5 | 79.10000000000001 |
| Paired Reads | 161870122 | 141132800 |
| Paired Reads (QC-failed) | 0 | 0 |
| Read1 | 80935061 | 70566400 |
| Read1 (QC-failed) | 0 | 0 |
| Read2 | 80935061 | 70566400 |
| Read2 (QC-failed) | 0 | 0 |
| Properly Paired Reads | 98226484 | 109823926 |
| Properly Paired Reads (QC-failed) | 0 | 0 |
| % Properly Paired Reads | 60.699999999999996 | 77.8 |
| With itself | 98678724 | 110336158 |
| With itself (QC-failed) | 0 | 0 |
| Singletons | 909003 | 1307324 |
| Singletons (QC-failed) | 0 | 0 |
| % Singleton | 0.6 | 0.8999999999999999 |
| Diff. Chrms | 16547 | 27794 |
| Diff. Chrms (QC-failed) | 0 | 0 |

### Marking duplicates (filtered BAM)

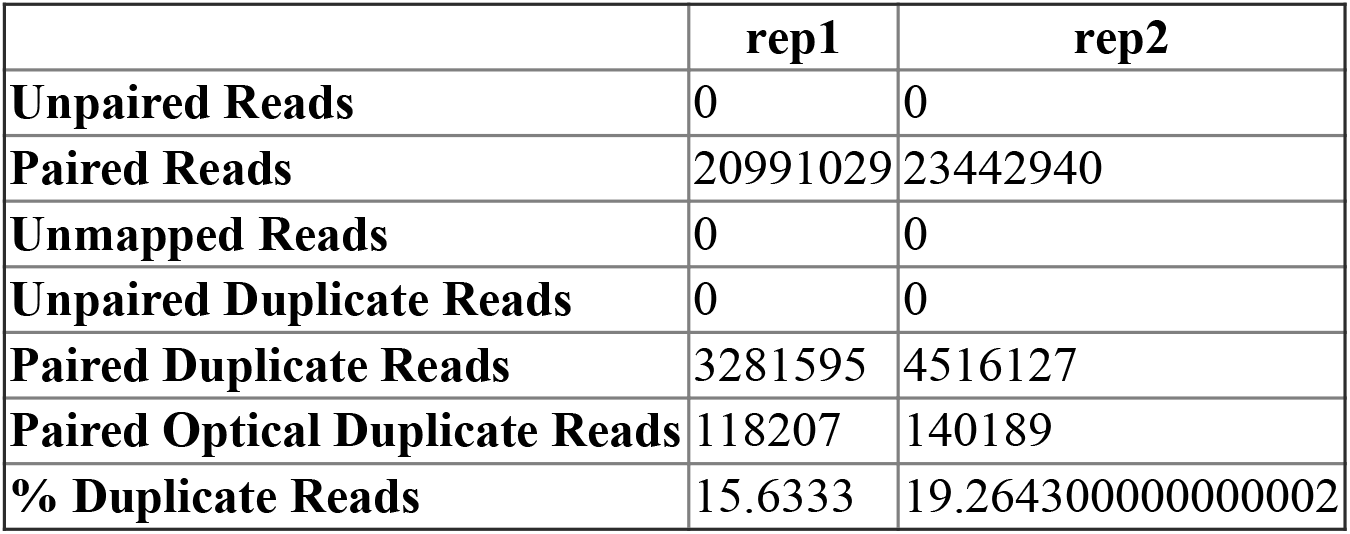

Filtered out (samtools view -F 1804):

- read unmapped (0×4)
- mate unmapped (0×8, for paired-end)
- not primary alignment (0×100)
- read fails platform/vendor quality checks (0×200)
- read is PCR or optical duplicate (0×400)

### Fraction of mitochondrial reads (unfiltered BAM)

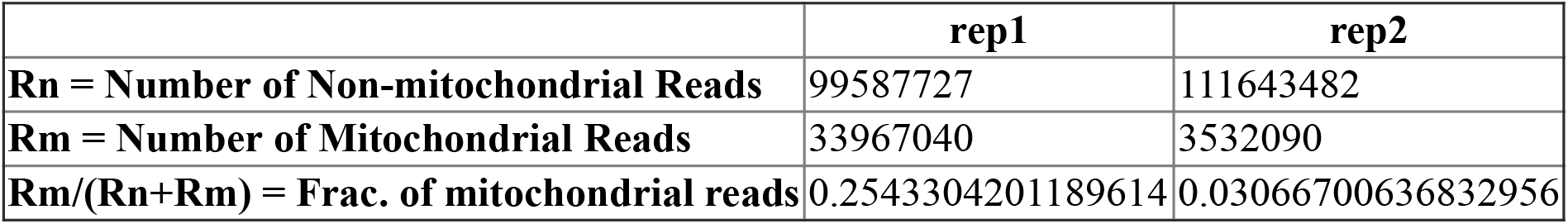

|  | rep1 | rep2 |
| --- | --- | --- |
| **Rn = Number of Non-mitochondrial Reads** | 99587727 | 111643482 |
| **Rm = Number of Mitochondrial Reads** | 33967040 | 3532090 |
| **Rm/(Rn+Rm) = Frac. of mitochondrial reads** | 0.2543304201189614 | 0.03066700636832956 |

### SAMstat (filtered/deduped BAM)

|  | rep1 | rep2 |
| --- | --- | --- |
| **Total Reads** | 35418868 | 37853626 |
| **Total Reads (QC-failed)** | 0 | 0 |
| **Duplicate Reads** | 0 | 0 |
| **Duplicate Reads (QC-failed)** | 0 | 0 |
| **Mapped Reads** | 35418868 | 37853626 |
| **Mapped Reads (QC-failed)** | 0 | 0 |
| **% Mapped Reads** | 100.0 | 100.0 |
| **Paired Reads** | 35418868 | 37853626 |
| **Paired Reads (QC-failed)** | 0 | 0 |
| **Read1** | 17709434 | 18926813 |
| **Read1 (QC-failed)** | 0 | 0 |
| **Read2** | 17709434 | 18926813 |
| **Read2 (QC-failed)** | 0 | 0 |
| **Properly Paired Reads** | 35418868 | 37853626 |
| **Properly Paired Reads (QC-failed)** | 0 | 0 |
| **% Properly Paired Reads** | 100.0 | 100.0 |
| **With itself** | 35418868 | 37853626 |
| **With itself (QC-failed)** | 0 | 0 |
| **Singletons** | 0 | 0 |
| **Singletons (QC-failed)** | 0 | 0 |
| **% Singleton** | 0.0 | 0.0 |
| **Diff. Chrms** | 0 | 0 |
| **Diff. Chrms (QC-failed)** | 0 | 0 |

Filtered and duplicates removed

### Fragment length statistics (filtered/deduped BAM)

|  | rep1 | rep2 |
| --- | --- | --- |
| **Fraction of reads in NFR** | 0.9323787083560022 | 0.8902688074183567 |
| **Fraction of reads in NFR (QC pass)** | True | True |
| **Fraction of reads in NFR (QC reason)** | OK | OK |
| **NFR / mono-nuc reads** | 15.598049844737124 | 9.061774693997215 |
| **NFR / mono-nuc reads (QC pass)** | True | True |
| **NFR / mono-nuc reads (QC reason)** | OK | OK |
| **Presence of NFR peak** | True | True |
| **Presence of Mono-Nuc peak** | False | True |
| **Presence of Di-Nuc peak** | False | False |

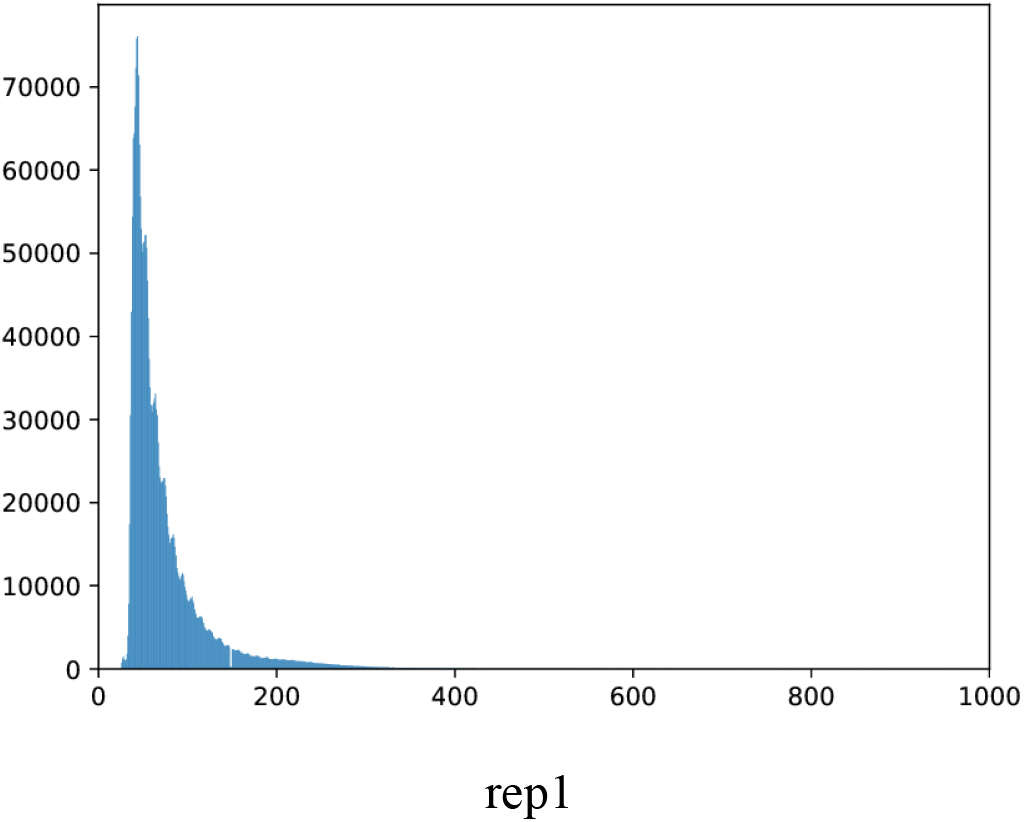
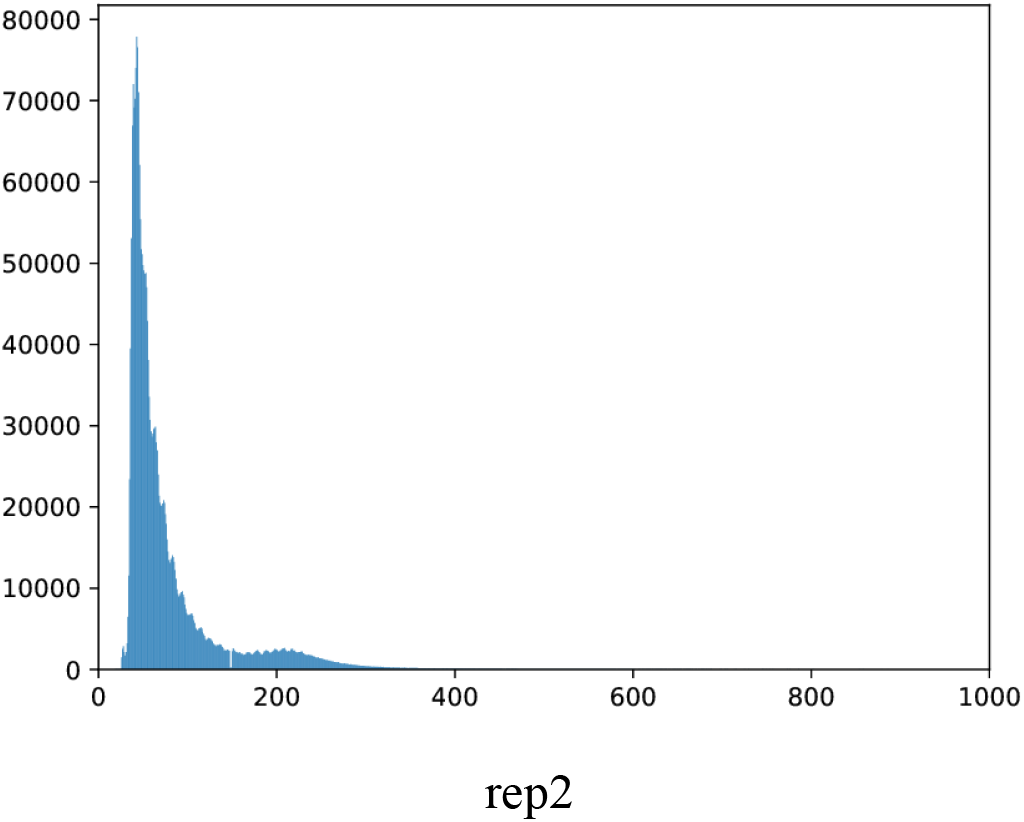

Open chromatin assays show distinct fragment length enrichments, as the cut sites are only in open chromatin and not in nucleosomes. As such, peaks representing different n-nucleosomal (ex mono-nucleosomal, di-nucleosomal) fragment lengths will arise. Good libraries will show these peaks in a fragment length distribution and will show specific peak ratios.

- NFR: Nucleosome free region

### Sequence quality metrics (filtered/deduped BAM)

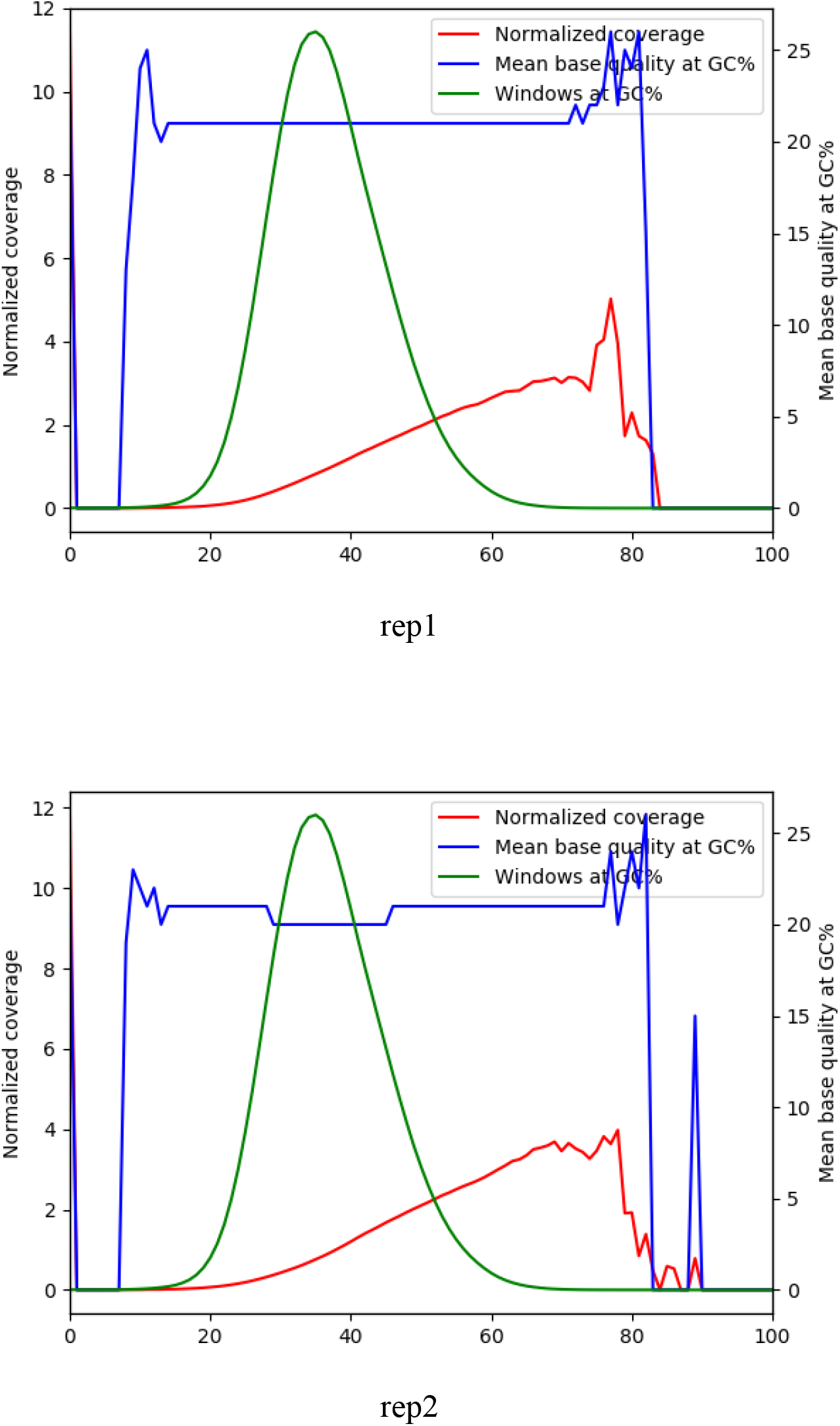

Open chromatin assays are known to have significant GC bias. Please take this into consideration as necessary.

## Library complexity quality metrics

### Library complexity (filtered non-mito BAM)

|  | rep1 | rep2 |
| --- | --- | --- |
| **Total Fragments** | 20991029 | 23442940 |
| **Distinct Fragments** | 17920070 | 19255358 |
| **Positions with Two Read** | 2196429 | 2734948 |
| **NRF = Distinct/Total** | 0.853701 | 0.821371 |
| **PBC1 = OneRead/Distinct** | 0.856182 | 0.825603 |
| **PBC2 = OneRead/TwoRead** | 6.985358 | 5.812643 |

Mitochondrial reads are filtered out by default. The non-redundant fraction (NRF) is the fraction of non-redundant mapped reads in a dataset; it is the ratio between the number of positions in the genome that uniquely mapped reads map to and the total number of uniquely mappable reads. The NRF should be > 0.8. The PBC1 is the ratio of genomic locations with EXACTLY one read pair over the genomic locations with AT LEAST one read pair. PBC1 is the primary measure, and the PBC1 should be close to 1. Provisionally 0-0.5 is severe bottlenecking, 0.5-0.8 is moderate bottlenecking, 0.8-0.9 is mild bottlenecking, and 0.9-1.0 is no bottlenecking. The PBC2 is the ratio of genomic locations with EXACTLY one read pair over the genomic locations with EXACTLY two read pairs. The PBC2 should be significantly greater than 1.

NRF (non redundant fraction)
PBC1 (PCR Bottleneck coefficient 1)
PBC2 (PCR Bottleneck coefficient 2)
PBC1 is the primary measure. Provisionally

- 0-0.5 is severe bottlenecking
- 0.5-0.8 is moderate bottlenecking
- 0.8-0.9 is mild bottlenecking
- 0.9-1.0 is no bottlenecking

## Replication quality metrics

### IDR (Irreproducible Discovery Rate) plots

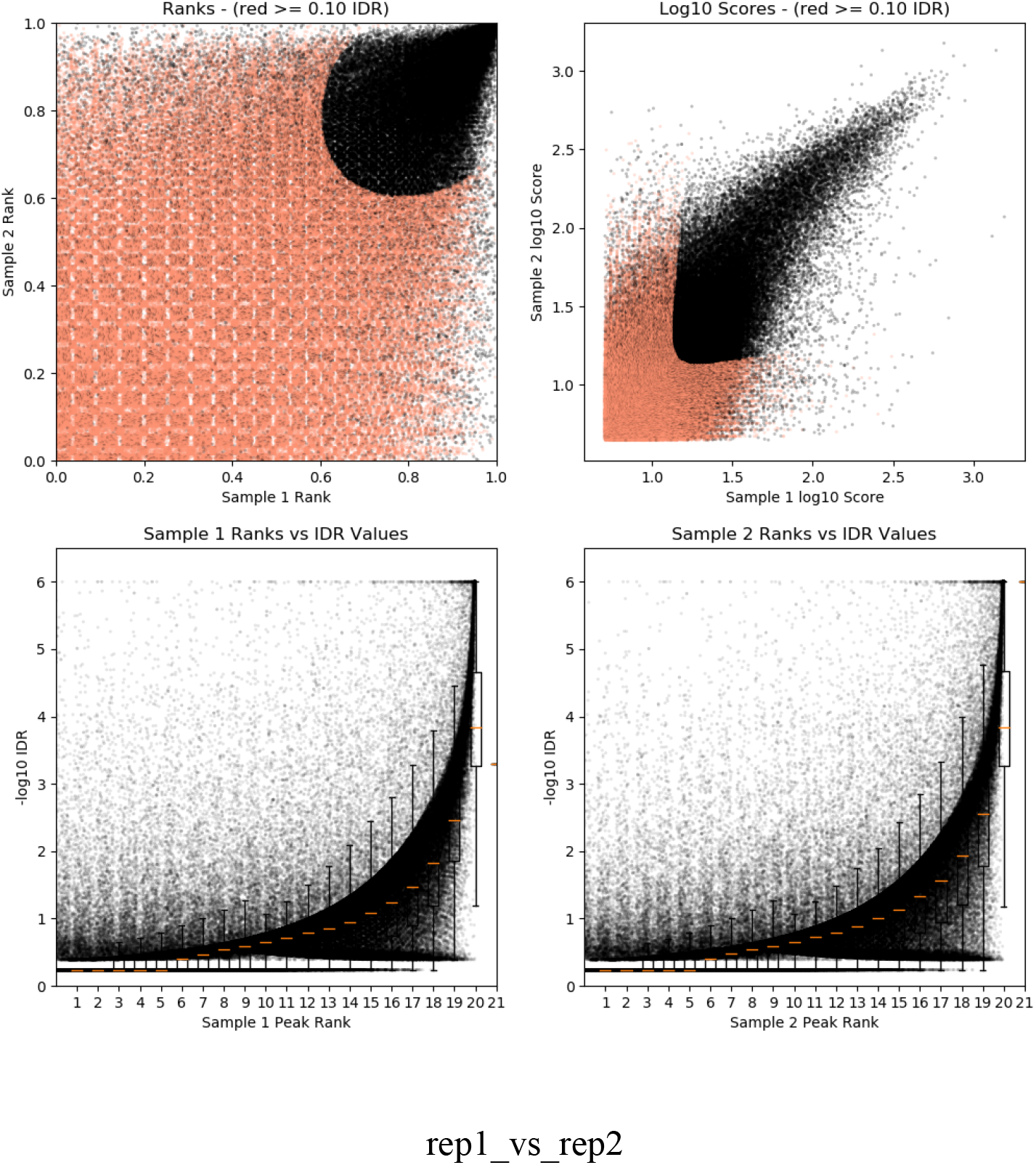

### Number of raw peaks

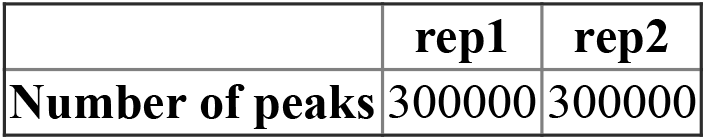

|  | rep1 | rep2 |
| --- | --- | --- |
| Number of peaks | 300000 | 300000 |

Top 300000 raw peaks from macs2 with p-val threshold 0.05

## Peak calling statistics

### Peak region size

|  | rep1 | rep2 |
| --- | --- | --- |
| Min size | 150.0 | 150.0 |
| 25 percentile | 223.0 | 223.0 |
| 50 percentile (median) | 348.0 | 355.0 |
| **75 percentile** | 567.0 | 582.0 |
| **Max size** | 3602.0 | 4282.0 |
| **Mean** | 453.15470666666664 | 461.8133433333333 |

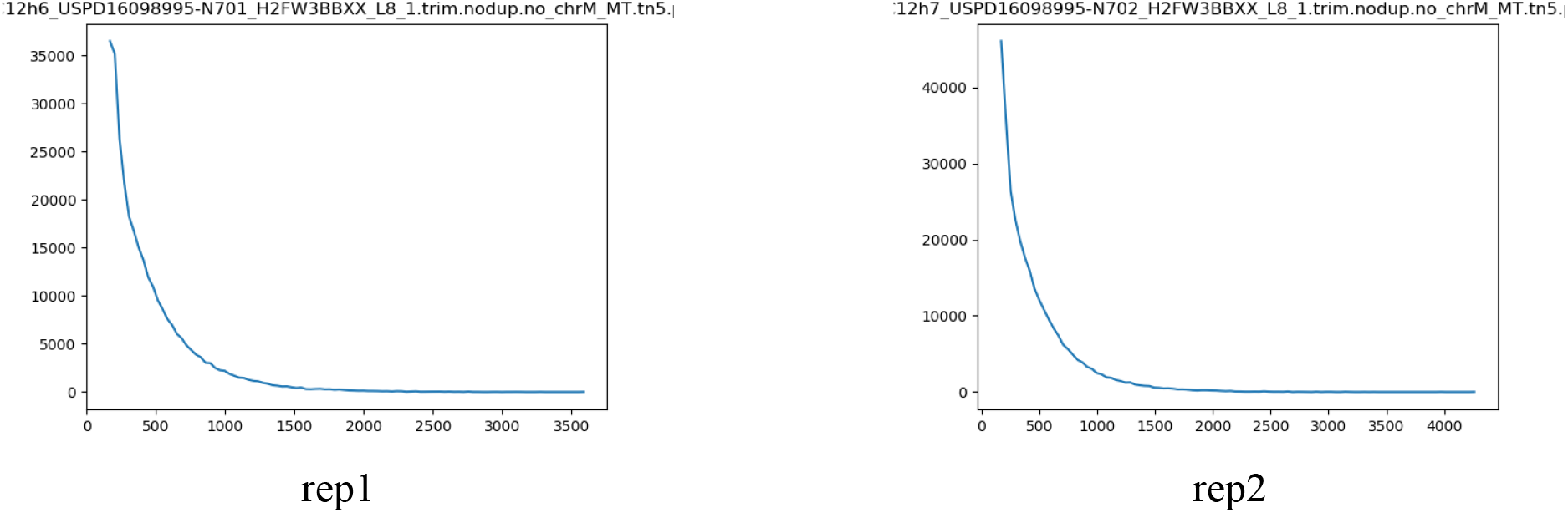

## Peak enrichment

### Fraction of reads in peaks (FRiP)

#### FRiP for macs2 raw peaks

|  | rep1 | rep2 | pooled |
| --- | --- | --- | --- |
| **Fraction of Reads in Peaks** | 0.5425538444650462 | 0.5892748821473536 | 0.5670562817201227 |

#### FRiP for overlap peaks

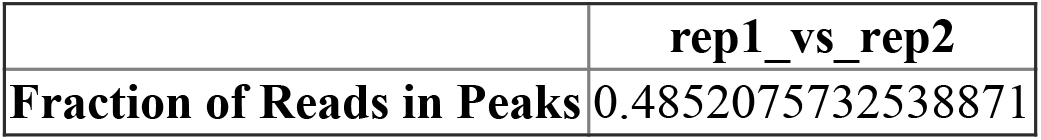

#### FRiP for IDR peaks

|  | rep1_vs_rep2 |
| --- | --- |
| **Fraction of Reads in Peaks** | 0.2598057328306581 |

For macs2 raw peaks:

- repX: Peak from true replicate X
- repX-prY: Peak from Yth pseudoreplicates from replicate X
- pooled: Peak from pooled true replicates (pool of rep1, rep2, …)
- pooled-pr1: Peak from 1st pooled pseudo replicate (pool of rep1-pr1, rep2-pr1, …)
- pooled-pr2: Peak from 2nd pooled pseudo replicate (pool of rep1-pr2, rep2-pr2, …)

For overlap/IDR peaks:

- repX_vs_repY: Comparing two peaks from true replicates X and Y
- repX-pr1_vs_repX-pr2: Comparing two peaks from both pseudoreplicates from replicate X
- pooled-pr1_vs_pooled-pr2: Comparing two peaks from 1st and 2nd pooled pseudo replicates

## QC Report

|  | general |
| --- | --- |
| Report generated at | 2020-04-22 02:18:15 |
| Title | ATAC_20h_3.1 |
| Description | PE and SR |
| Pipeline version | v1.7.0 |
| Pipeline type | atac |
| Genome | Sp3.1 |
| Aligner | bowtie2 |
| Sequencing endedness | {'rep1': {'paired_end': True}, 'rep2': {'paired_end': False}, 'rep3': {'paired_end': False}} |
| Peak caller | macs2 |

## Alignment quality metrics

### SAMstat (raw unfiltered BAM)

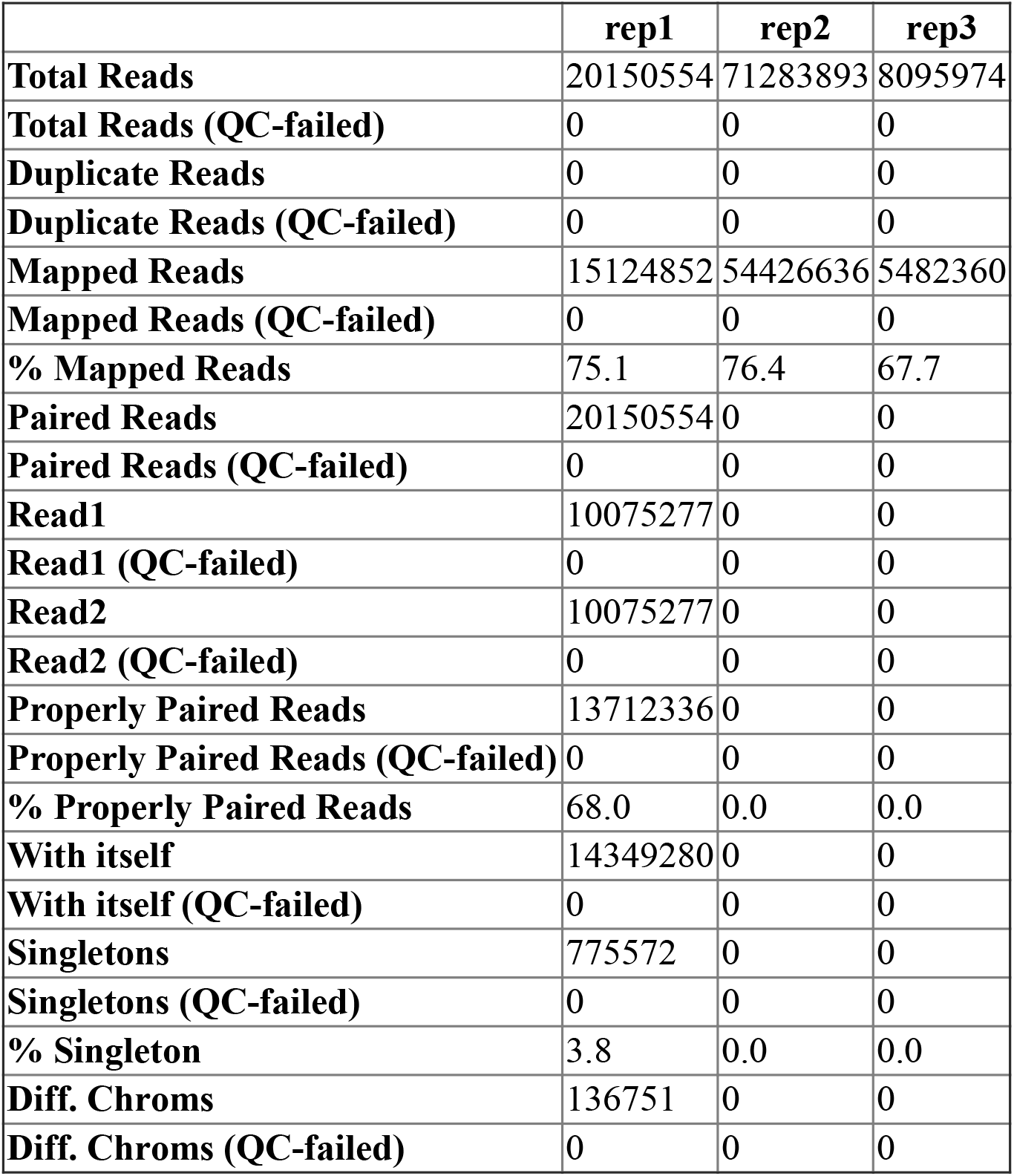

Alignment quality metrics SAMstat (raw unfiltered BAM)
|  | rep1 | rep2 | rep3 |
| --- | --- | --- | --- |
| Total Reads | 20150554 | 71283893 | 8095974 |
| Total Reads (QC-failed) | 0 | 0 | 0 |
| Duplicate Reads | 0 | 0 | 0 |
| Duplicate Reads (QC-failed) | 0 | 0 | 0 |
| Mapped Reads | 15124852 | 54426636 | 5482360 |
| Mapped Reads (QC-failed) | 0 | 0 | 0 |
| % Mapped Reads | 75.1 | 76.4 | 67.7 |
| Paired Reads | 20150554 | 0 | 0 |
| Paired Reads (QC-failed) | 0 | 0 | 0 |
| Read1 | 10075277 | 0 | 0 |
| Read1 (QC-failed) | 0 | 0 | 0 |
| Read2 | 10075277 | 0 | 0 |
| Read2 (QC-failed) | 0 | 0 | 0 |
| Properly Paired Reads | 13712336 | 0 | 0 |
| Properly Paired Reads (QC-failed) | 0 | 0 | 0 |
| % Properly Paired Reads | 68.0 | 0.0 | 0.0 |
| With itself | 14349280 | 0 | 0 |
| With itself (QC-failed) | 0 | 0 | 0 |
| Singletons | 775572 | 0 | 0 |
| Singletons (QC-failed) | 0 | 0 | 0 |
| % Singleton | 3.8 | 0.0 | 0.0 |
| Diff. Chroms | 136751 | 0 | 0 |
| Diff. Chroms (QC-failed) | 0 | 0 | 0 |

### Marking duplicates (filtered BAM)

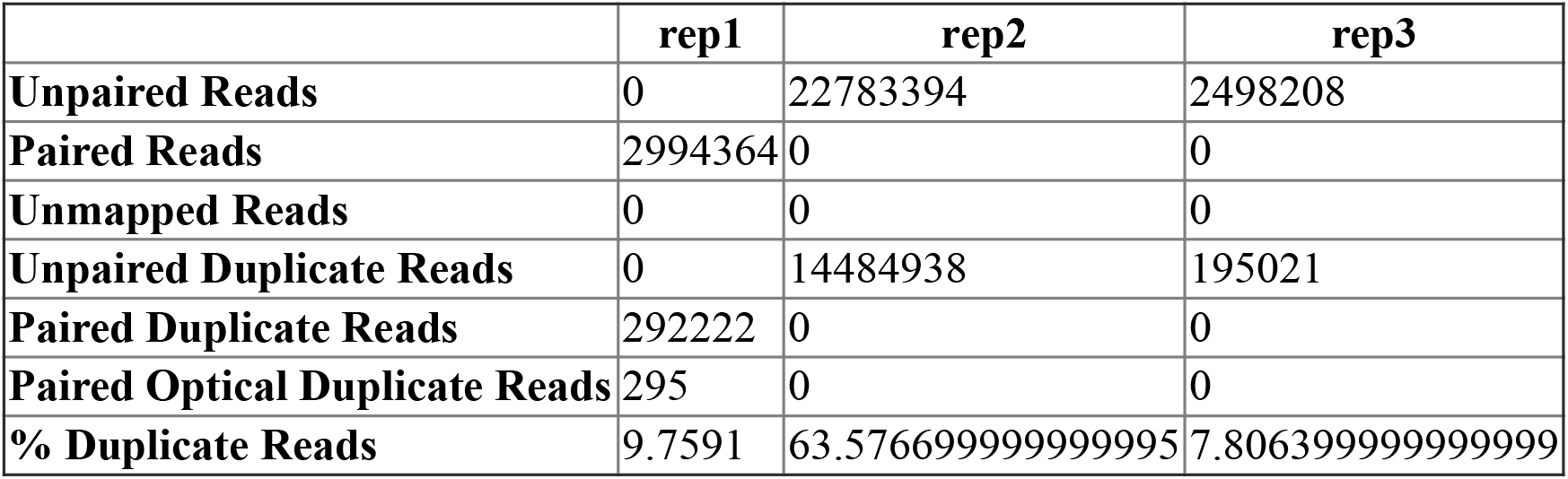

|  | rep1 | rep2 | rep3 |
| --- | --- | --- | --- |
| **Unpaired Reads** | 0 | 22783394 | 2498208 |
| **Paired Reads** | 2994364 | 0 | 0 |
| **Unmapped Reads** | 0 | 0 | 0 |
| **Unpaired Duplicate Reads** | 0 | 14484938 | 195021 |
| **Paired Duplicate Reads** | 292222 | 0 | 0 |
| **Paired Optical Duplicate Reads** | 295 | 0 | 0 |
| **% Duplicate Reads** | 9.7591 | 63.576699999999995 | 7.806399999999999 |

Filtered out (samtools view -F 1804):

### Fraction of mitochondrial reads (unfiltered BAM)

|  | rep1 | rep2 | rep3 |
| --- | --- | --- | --- |
| **Rn = Number of Non-mitochondrial Reads** | 15124852 | 54426636 | 5482360 |
| **Rm = Number of Mitochondrial Reads** | 153339 | 589223 | 1311787 |
| **Rm/(Rn+Rm) = Frac. of mitochondrial reads** | 0.010036463086500228 | 0.010710057258217127 | 0.1930760403035142 |

### SAMstat (filtered/deduped BAM)

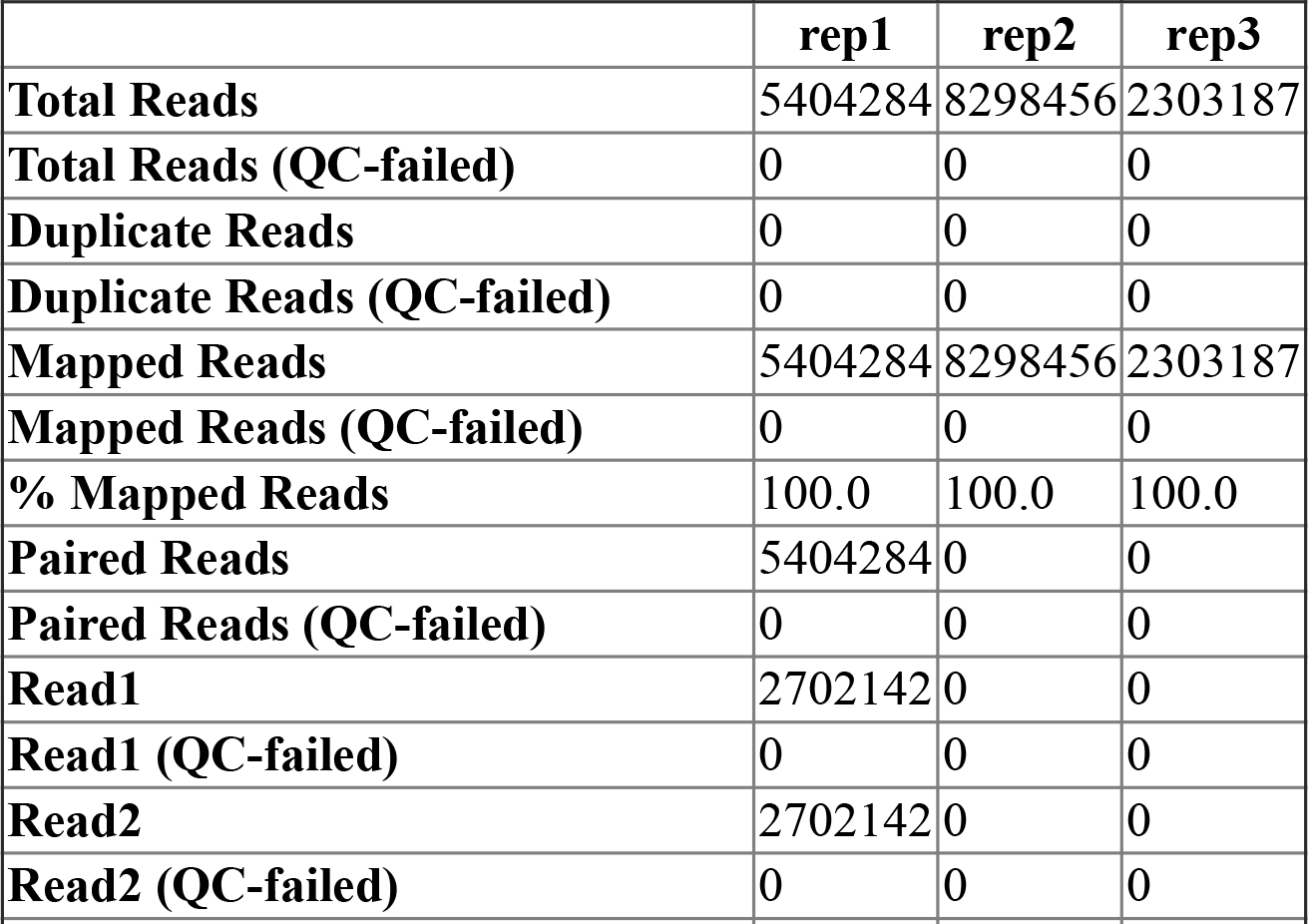
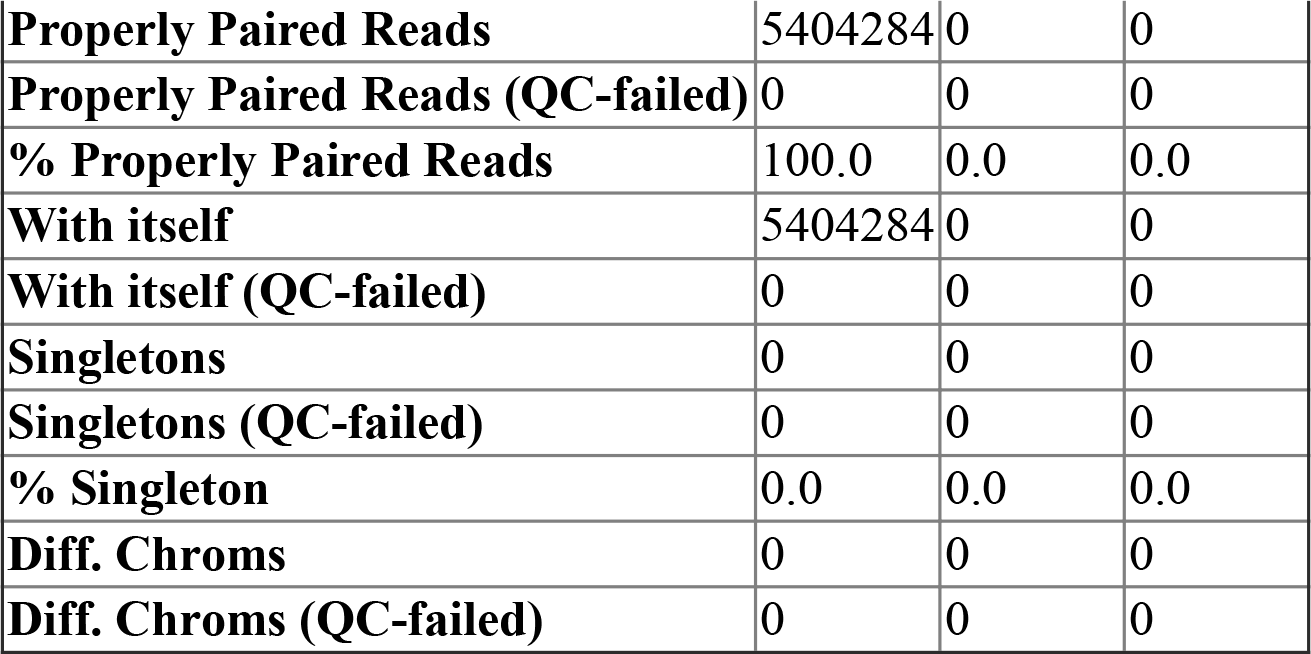

Filtered and duplicates removed

### Fragment length statistics (filtered/deduped BAM)

|  |  |
| --- | --- |
|  | rep1 |
| Fraction of reads in NFR | 0.6477683030192886 |
| Fraction of reads in NFR (QC pass) | True |
| Fraction of reads in NFR (QC reason) | OK |
| NFR / mono-nuc reads | 2.733468616968029 |
| NFR / mono-nuc reads (QC pass) | True |
| NFR / mono-nuc reads (QC reason) | OK |
| Presence of NFR peak | True |
| Presence of Mono-Nuc peak | True |
| Presence of Di-Nuc peak | True |

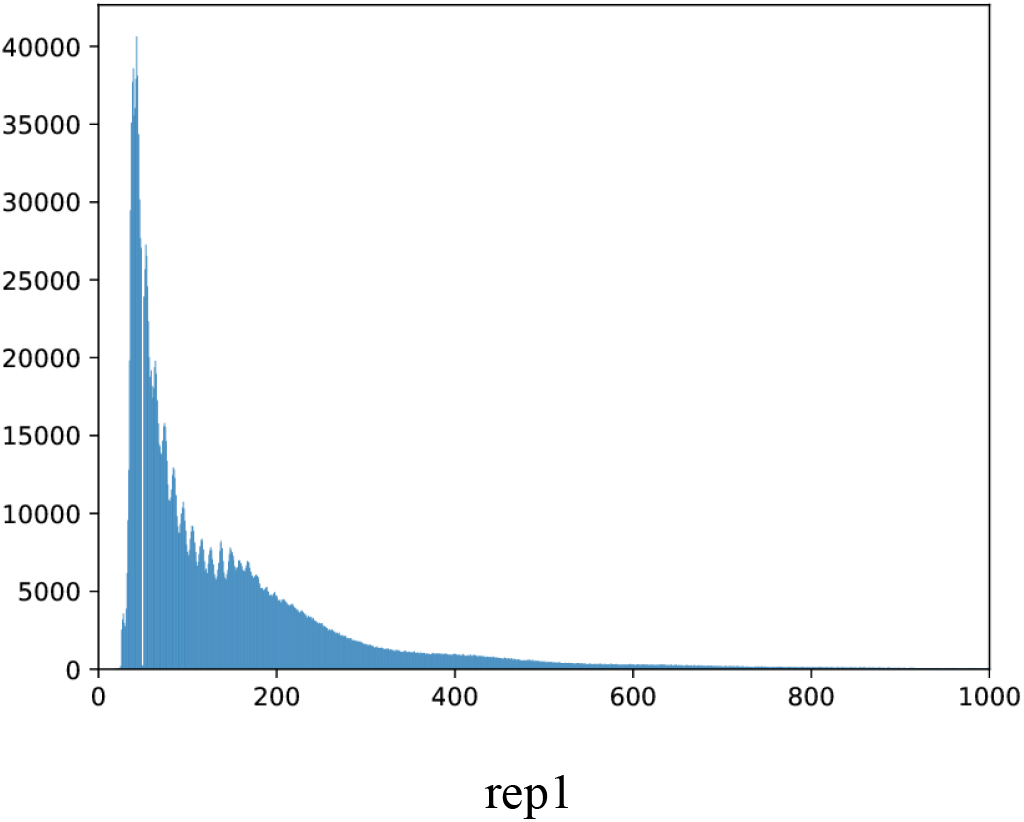

- NFR: Nucleosome free region

### Sequence quality metrics (filtered/deduped BAM)

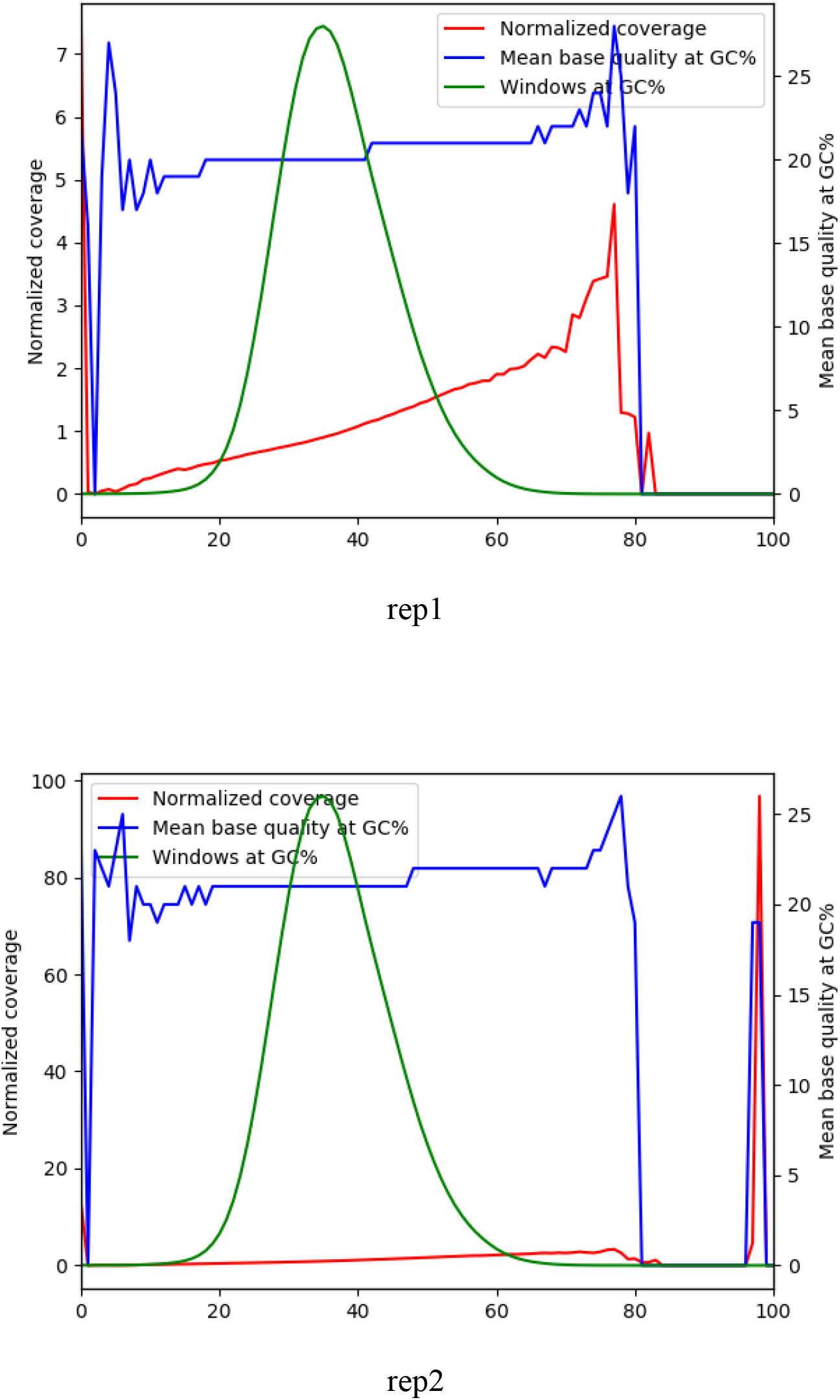
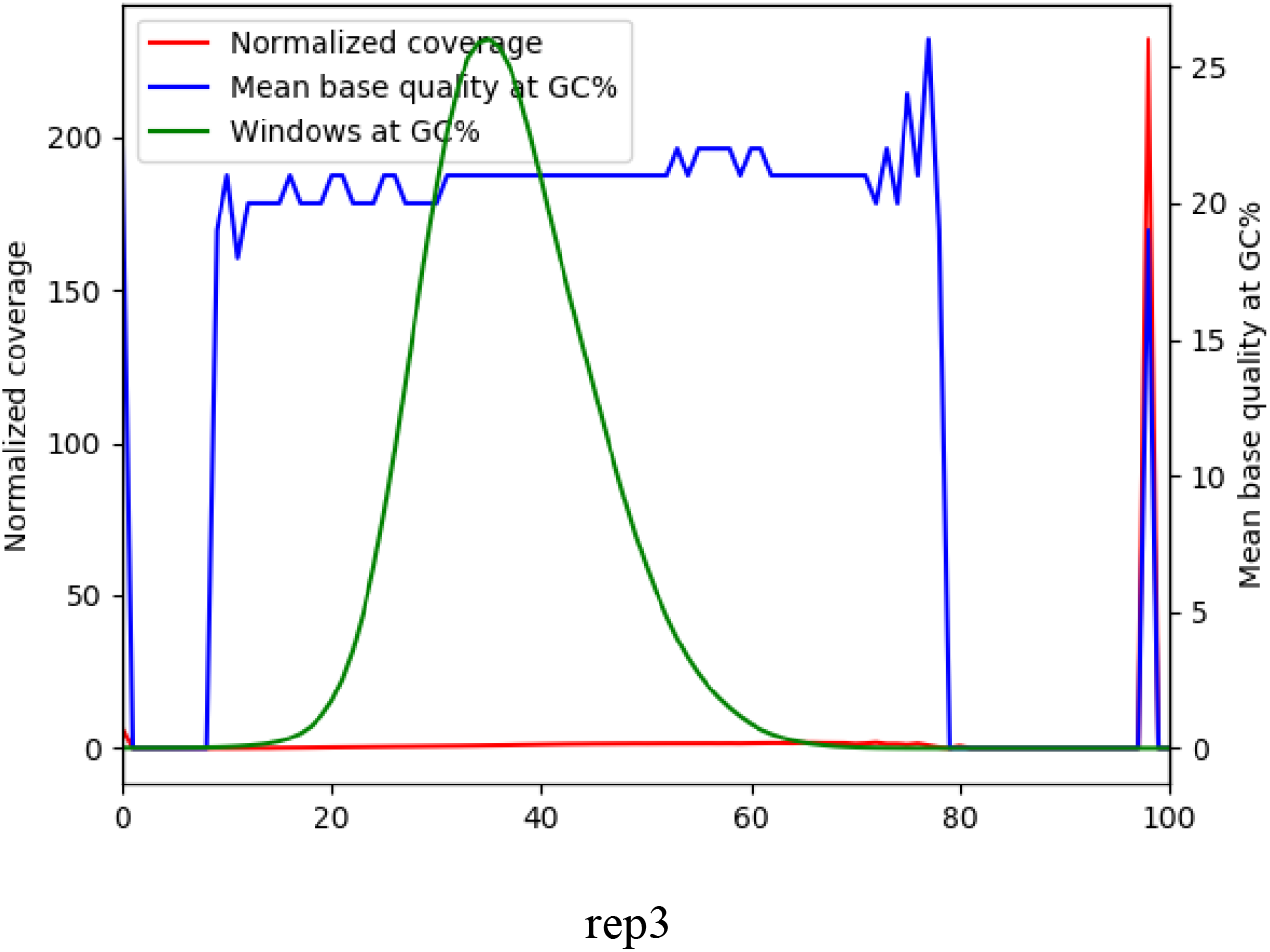

## Library complexity quality metrics

### Library complexity (filtered non-mito BAM)

|  | rep1 | rep2 | rep3 |
| --- | --- | --- | --- |
| **Total Fragments** | 2994364 | 22783394 | 2498208 |
| **Distinct Fragments** | 2702613 | 8533758 | 2335470 |
| **Positions with Two Read** | 243447 | 2119342 | 119191 |
| **NRF = Distinct/Total** | 0.902567 | 0.37456 | 0.934858 |
| **PBC1 = OneRead/Distinct** | 0.901346 | 0.318471 | 0.941364 |
| **PBC2 = OneRead/TwoRead** | 10.00624 | 1.282357 | 18.44542 |

## Replication quality metrics

### IDR (Irreproducible Discovery Rate) plots

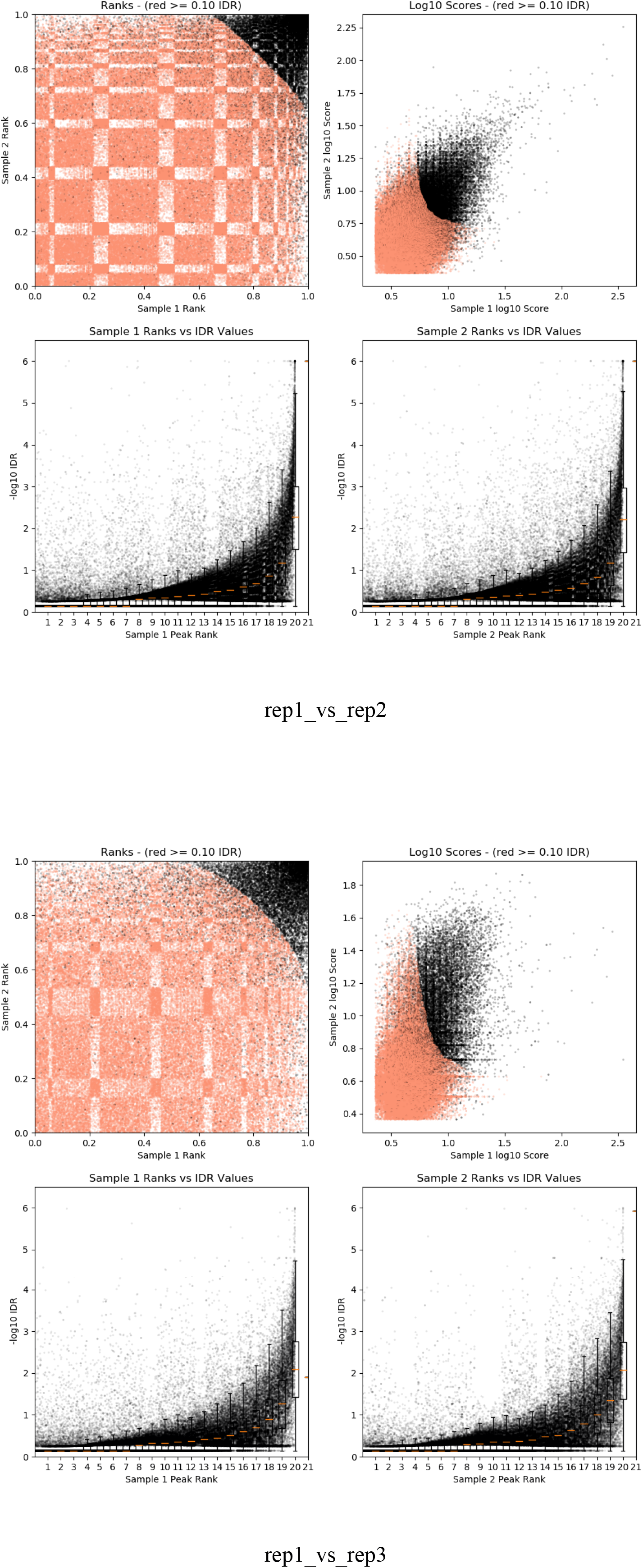
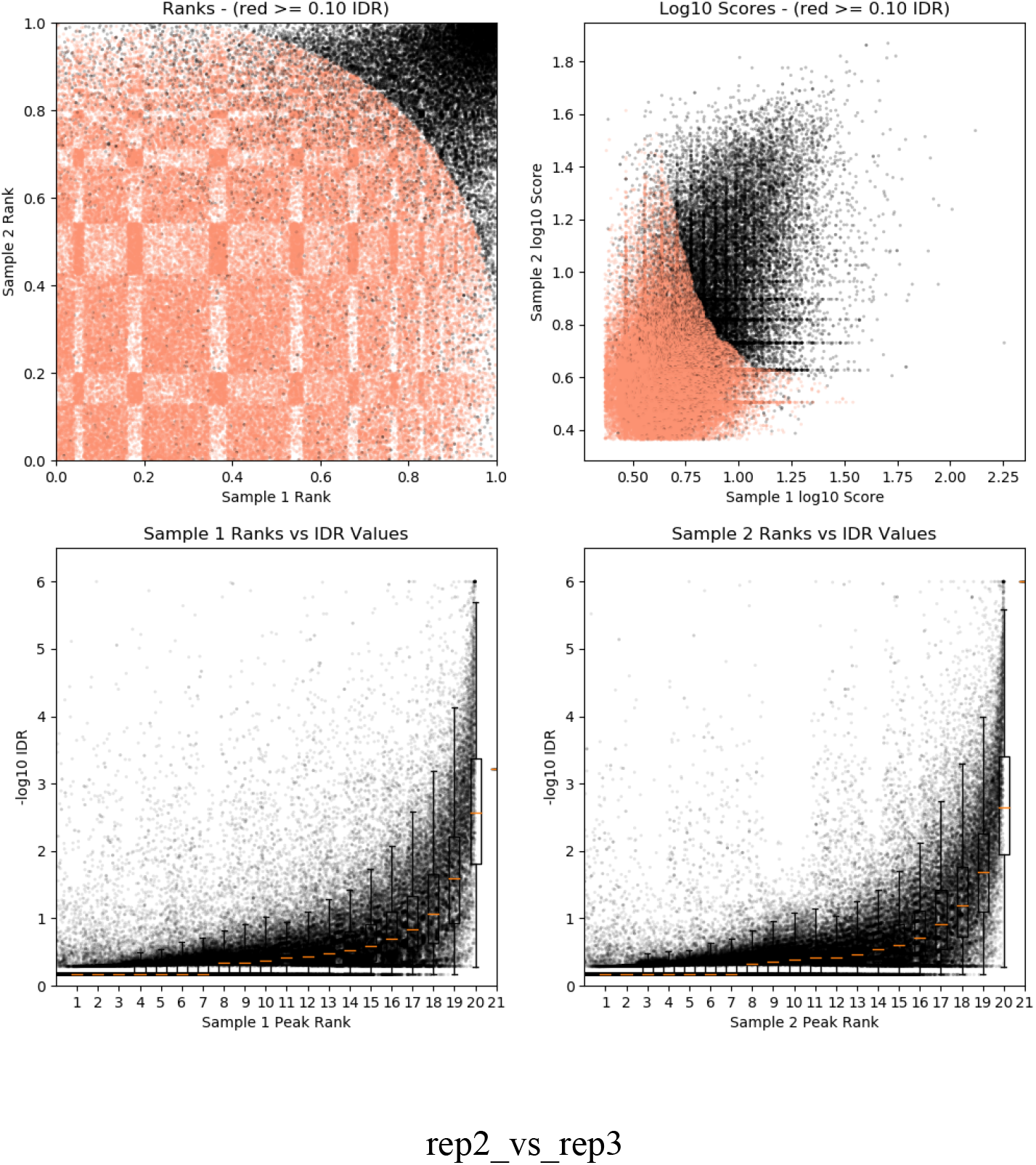

### Number of raw peaks

|  | rep1 | rep2 | rep3 |
| --- | --- | --- | --- |
| Number of peaks | 248700 | 300000 | 160654 |

Top 300000 raw peaks from macs2 with p-val threshold 0.05

## Peak calling statistics

### Peak region size

|  | rep1 | rep2 | rep3 |
| --- | --- | --- | --- |
| Min size | 150.0 | 150.0 | 150.0 |
| 25 percentile | 157.0 | 166.0 | 150.0 |
| 50 percentile (median) | 212.0 | 230.0 | 182.0 |
| 75 percentile | 313.0 | 353.0 | 299.0 |
| Max size | 3115.0 | 4044.0 | 2008.0 |
| Mean | 270.7728266988339 | 302.13554666666664 | 266.0243317937929 |

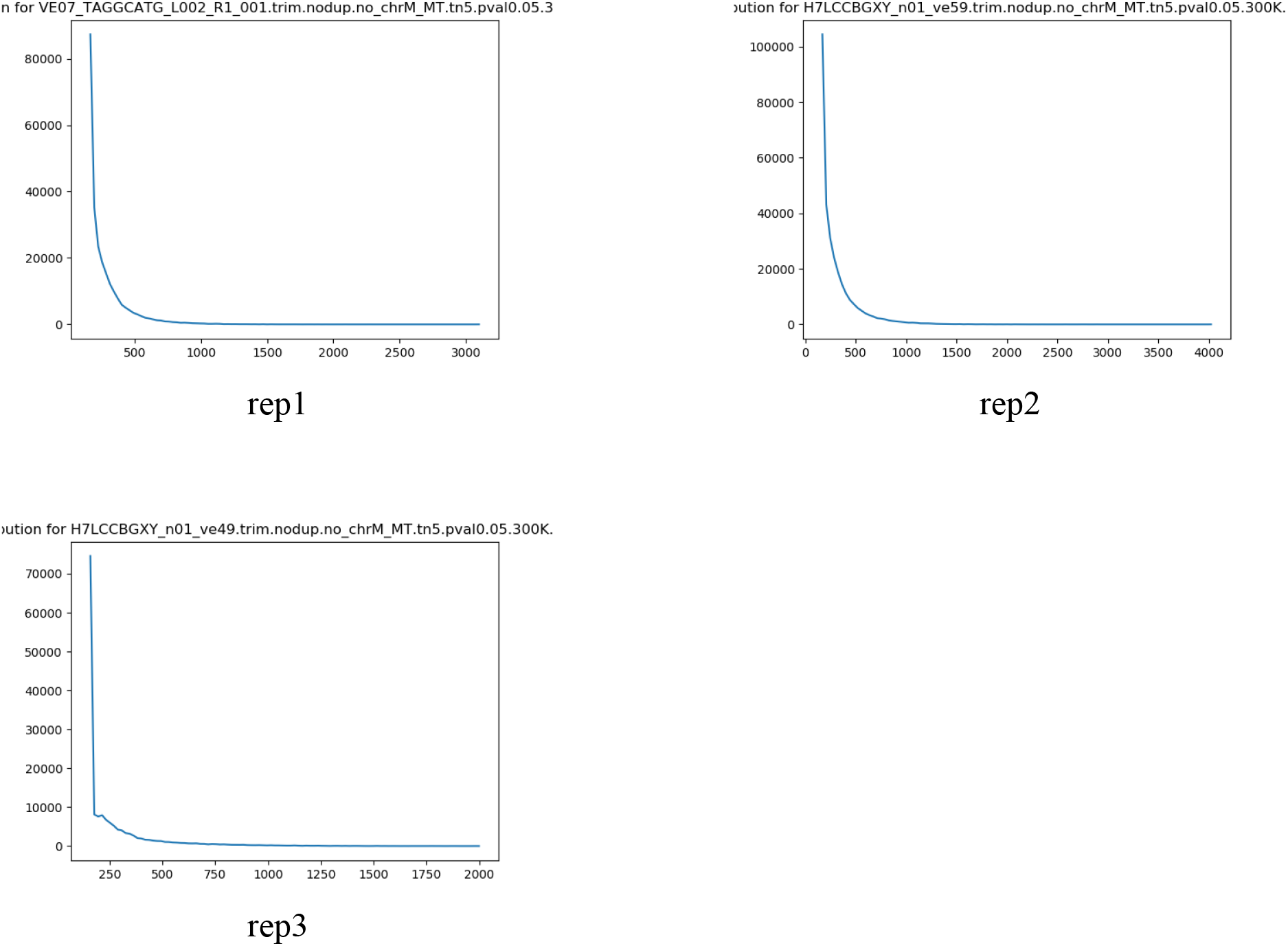

## Peak enrichment

### Fraction of reads in peaks (FRiP)

#### FRiP for macs2 raw peaks

|  | rep1 | rep2 | rep3 | pooled |
| --- | --- | --- | --- | --- |
| **Fraction of Reads in Peaks** | 0.4482899122251902 | 0.44582546439964255 | 0.495024937184866 | 0.44154006200328166 |

#### FRiP for overlap peaks

|  | rep1_vs_rep2 | rep1_vs_rep3 | rep2_vs_rep3 |
| --- | --- | --- | --- |
| **Fraction of Reads in Peaks** | 0.29597523467400544 | 0.1967762317046679 | 0.21487827602862364 |

#### FRiP for IDR peaks

|  | rep1_vs_rep2 | rep1_vs_rep3 | rep2_vs_rep3 |
| --- | --- | --- | --- |
| **Fraction of Reads in Peaks** | 0.07651753003746675 | 0.05970275885926507 | 0.07640663361765926 |

For macs2 raw peaks:

For overlap/IDR peaks:

